# Genetic Dissection of Heat Stress Tolerance in Soybean through Genome-Wide Association Studies and the Use of Genomic Prediction to Enhance Breeding Applications

**DOI:** 10.1101/2024.04.27.591454

**Authors:** Liza Van der Laan, Leonardo de Azevedo Peixoto, Asheesh K. Singh

## Abstract

Rising temperatures and associated heat stress pose an increasing threat to soybean [*Glycine max* L. (Merr.)] productivity. Due to a limited choice of mitigation strategies, the primary arsenal in crop protection comes from improved genetic stress tolerance. Despite this current and looming threat to soybean production, limited studies have examined the genetics of heat stress tolerance. There is a need to conduct large-scale germplasm screening and genetic studies, including genome-wide association mapping and genomic prediction, to identify genomic regions and useful markers associated with heat tolerance traits that can be utilized in soybean breeding programs. We screened a diverse panel of 450 soybean accessions from MG 0-IV to dissect the genetic architecture of physiological and growth-related traits under optimal and heat stress temperatures and study trait relationships and predictive ability. The genetic architecture information of the response to heat revealed in this study provides insights into the genetics of heat stress tolerance. Thirty-seven significant SNPs were detected, with 20 unique SNPs detected in optimal, 16 detected in heat stress, and a single SNP detected for a heat tolerance index. Only one significant SNP was identified across temperature treatments indicating a genetic divergence in soybean responses to temperature. The genomic prediction worked well for biomass traits, but physiological traits associated with heat stress had poor model accuracy. Through our phenotyping efforts, we identified heat tolerant soybean accessions. The identification of heat tolerant accessions and significant SNPs are useful in heat tolerant variety development through marker-assisted and genomic selection.

**Core ideas:**

- Soybean exhibit phenotypic diversity in response to heat stress.
- Large scale phenotypic screening identified heat tolerant accessions.
- Previously unreported QTL and SNP associated with biomass and physiological parameters under heat stress are reported.
- Genomic prediction shows promise in abiotic stress breeding applications.

## 1 INTRODUCTION

Soybean [*Glycine max* L. (Merr.)] is a major oilseed crop (OECD and FAO - United Nations, 2023) belonging to the Leguminosae family, with origins in the temperate regions of China ∼5000 years ago (Sedivy et al., 2017). Soybean acres have rapidly expanded globally and span a wide latitude distribution (Leff et al., 2004), demonstrating its adaptability to different environments. Soybean, similar to other row crops, is adversely affected by abiotic stresses, which, individually or in combination, cause changes that negatively affect crop growth and productivity. Soybean are a warm-season legume with a C3 system of photosynthesis (Vu et al., 1997) and are considered a relatively heat-tolerant crop in comparison to cool-season legume crops such as field pea (*Pisum sativum* L.). However, heat, drought, cold, and salinity are the primary climatic abiotic factors that cause severe crop damage and associated yield losses (Oerke, 2006).

Climate change, including elevated CO2 levels, will impact soybean agriculture, though any yield increases from CO2 fertilization are negated by drought and heat stresses (Hatfield et al., 2011); (Gray et al., 2016; Jin et al., 2018; Thomey et al., 2019)). The long term forecast of changing climatic conditions including increase in temperatures threatens all crops, including legume crops (Teixeira et al., 2013; Mousavi-Derazmahalleh et al., 2019; Sun et al., 2019; IPCC, 2023; Yang and Wang, 2023). Plants have an inherent and evolved mechanism to respond to abiotic stresses, which are considered a complex trait since it involves numerous metabolic pathways, cellular, and molecular components (Singh et al., 2021). Plant response to abiotic stresses involves mechanisms that are interconnected, leading to cellular damage and secondary stresses, including osmotic and oxidative stresses. The inheritance of abiotic stress tolerance is multi-genic, generally with smaller effects controlling the trait (Deshmukh et al., 2014).

The effect of heat stress on crop morphology, physiology, and reproduction has been well-studied in various crop species (Bita and Gerats, 2013; Prasad et al., 2017; Jagadish et al., 2021), although the responses differ across species and genotypes. Heat imposed in vegetative tissue has been found to negatively affect vital physiological processes, including chlorophyll content, photosynthesis, cellular respiration, and stomatal conductance (Almeselmani et al., 2012; Moore et al., 2021; Poudel et al., 2023). Additionally, heat stress has been found to increase Reactive Oxygen Species (ROS) activity, which affects the cellular structure and the production of pigments and can deteriorate the thylakoid membranes (Jumrani and Bhatia, 2019). These effects also occur in the plant during reproductive stages, which can be more at risk to heat due to the sensitivity of gametogenesis (Bita and Gerats, 2013), leading to crop yield losses due to fewer fertilization successes. Genetic variation of these traits has been used to distinguish sensitive and tolerant genotypes in various crops (Tesfaye et al., 2017; Balla et al., 2019), which can be utilized to breed for improved crop performance under stressful environments. Genetic studies aimed at exploring molecular mechanisms of heat stress response complement breeding efforts. These studies establish the mechanisms of heat stress tolerance and crop response; however, limited information is available for soybean and most of the heat-related studies come from other crop species, particularly in cereal species. Given the complexity of tolerance to abiotic stresses such as heat (Singh et al., 2021), breeding for heat tolerance is one of the most economical approaches for protecting soybean yields. Due to the paucity of data on heat stress response in soybean, large accession panels are required to sufficiently sample the genetic variation and study the genetics of heat tolerance.

In non-soybean legumes, Quantitative Trait Loci (QTL) and candidate genes have been reported for field peas (Tafesse et al., 2020), common bean (*Phaseolus vulgaris* L.) (López-Hernández and Cortés, 2019) peanuts (*Arachis hypogaea* L.) (Sharma et al., 2023), faba beans (*Vicia faba* L.) (Maalouf et al., 2022), and chickpeas (*Cicer arietinum* L.) (Paul et al., 2018) grown under heat stress. Sequencing studies of soybeans grown in heat conditions have shown an overexpression of heat shock proteins and transcription factors (Li et al., 2014; Valdés-López et al., 2016; Song et al., 2017b), as well as differentially expressed genes related to metabolic processes and molecular transports (Wang et al., 2018). Heat stress has been found to induce overexpression of the DREB1 gene family, LEA genes, and dehydrins, which were also overexpressed for other abiotic stresses (Kidokoro et al., 2015).

The response of soybean to heat stress is less studied than heat stress in cool season legume species, although soybean yield losses due to heat have been reported in numerous studies (Ruiz-Vera et al., 2013; Tacarindua et al., 2013; Siebers et al., 2015; Thomey et al., 2019; Alsajri et al., 2020; Cohen et al., 2021). Additionally, simulations of soybean yield at end-of-century conditions have found yield losses of up to 22% due to the effects of high temperatures (Yang and Wang, 2023), or a decline of 3.3% for each decade, with temperature rise being one of the main factors for the decline (IPCC, 2023). However, information on the impact of heat stress on early-season soybean growth is limited (Alsajri et al., 2019), as well as the genetic architecture that may be underlying soybean tolerance to heat stress is poorly understood. To date, no study has reported on GWAS detected genes associated with physiological traits of soybeans grown under heat stress conditions. Soybean has a wide genetic variation for multiple traits, as demonstrated in studies for yield and abiotic stress tolerance that identified candidate genes via GWAS for yield, seed composition (Zhang et al., 2018), drought (Kaler et al., 2017; Steketee et al., 2020), flooding (Wu et al., 2019), and iron deficiency chlorosis (Mamidi et al., 2011; Assefa et al., 2020).

Soybean growth and development stages include vegetative and reproductive phases, and previous research has identified that crop losses start when the average daily air temperature exceeds ∼ 28-30°C during the vegetative stage (Hesketh et al., 1973), and 22-24°C during the reproductive stage (Hatfield et al., 2011). However, these optimum temperatures for healthy plant development were proposed before the more dramatic shift in climate patterns and increasing temperatures. These temperature increases invariably coincide with water-limited conditions, i.e., drought. Drought has long been known as a major limiting factor in soybean productivity and is expected to remain a major limiting factor. However, it is predicted that a 2°C rise in global temperatures may cause heat to become the foremost cause of yield losses in soybean and maize in the US Corn Belt (Yang and Wang, 2023). As temperatures rise and water availability to plants at critical times becomes a dual threat of exceeding proportions, global soybean production will be severely impacted.

Although difficult and often uneconomical or unsustainable, irrigation can be provided to counter the water deficit. However, against heat, improved genetic materials currently remain the best solution. For example, one tool to counteract the adverse effects of future climates is to plant varieties with a genetic tolerance to various abiotic stresses, such as heat. A difficulty of developing soybean genotypes with superior tolerance to abiotic stress, such as heat, is that such tolerance genes are generally quantitative and are controlled by many small-effect QTLs. Plant breeders can use genomic tools, including marker assisted selection (MAS), genomic selection (GS) and genetic engineering to breed for soybean heat stress tolerance more rapidly. With genomic prediction, breeders can evaluate the genetic potential of a large number of genotypes without needing to grow and evaluate each genotype, which saves time, resources, and labor (Wang et al., 2020).

The objectives of this study were: (a) screen a diverse accession panel to examine the early season heat tolerance response in soybean, (b) understand the impact of heat on physiological traits with potential for HTP applications, (c) conduct genome-wide association studies to elucidate genomic regions crucial for heat stress tolerance and identify candidate genes for heat stress tolerance, and (d) investigate the usefulness of genomic prediction of different heat stress related physiological traits, and project the GP model to screening the USDA core collection across all maturity groups.

## 2 MATERIALS AND METHODS

### 2.1 Plant Materials

A panel of 450 diverse soybean accessions (de Azevedo Peixoto et al., 2017; Moellers et al., 2017) was used to study heat response in controlled environments (Table S1). A sub-set of 418 plant introduction (PI) lines were obtained from the USDA GRIN soybean mini-core collection (Oliveira et al., 2010). These PI lines originated from 26 countries. An additional 22 SoyNAM parents (Song et al., 2017a) and ten elite lines commonly used as checks in the Uniform Soybean Trials (UST) were selected for this study. A prior pre-screen process of the PIs was utilized to ensure that only accessions with good standability and non-shattering were included in this study. All accessions were selected for maturity suitable for production in Iowa and range in maturity groups 0-IV.

### 2.2 Experimental Design and Planting

The experimental design in each greenhouse was a randomized complete block with four replications in each temperature treatment. The replication was used as the blocking factor for placement within the greenhouse, with all four replications of a single treatment in the same greenhouse. To complete phenotypic measurements in a short amount of time to reduce for unaccounted variation from time lag, we planted each replication in the two treatments a day apart in a staggered manner, allowing complete phenotyping of a single replication of a single treatment in a day. For example, the first replication was planted in the control greenhouse on day 1. On day 2, the first replication was planted in the greenhouse with heat treatment. This sequence was followed for the entire experiment, allowing us to complete consistency in days after planting for data collection. The complete details of the days of planting and data collection are provided in Table S2.

Four seeds per accession were sown in a single Conetainer (Stuewe & Sons, Inc. Tangent, OR, USA) filled with sand. Each conetainer had a drainage hole that was covered with landscape fabric (Sta-Green Premium Landscape Fabric, Lowes) to prevent sand loss and ensure well-drained condition in the cones. A tablespoon of Osmocote slow-release fertilizer (14-14-14) was applied to each cone during planting. Cones were placed into their assigned temperature treatment at planting and remained in the designated treatment of the optimal control or the heat treatment until phenotyping. After emergence, plants were thinned to a single plant per cone. Plants were watered twice daily to ensure no drought stress would occur during the growing period.

### 2.3 Temperature Treatments

The experiment was conducted in two greenhouse rooms, one each for the optimal control (29/18°C day/night) and heat (36/29°C day/night) treatment greenhouses. To minimize all other sources of variance except the temperature treatments, we carefully monitored air, light, and temperature through regular monitoring.

### 2.4 Phenotypic Measurements

All phenotypic measurements were taken 28 days after planting. Measurements were taken on the center leaflet of the most recently fully developed trifoliate of each plant. Chlorophyll concentration (SPAD) was estimated non-destructively using a SPAD chlorophyll meter (Konica Minolta Sensing Americas Inc., USA). Chlorophyll concentration was measured as a SPAD value, which is unitless, and is calculated as the ratio of the intensity of light transmittance from 650-940 nm (red to infrared). Canopy temperature (CT) was measured using an infrared radiometer (Model SI-411-SS, Apogee Instruments, Logan, UT, USA) from 10 am to 2 pm to measure around the solar noon window. Biomass was collected from each plant and partitioned between shoot and root biomass. Samples were dried at 80°C for three days to remove all moisture and then weighed to get the dry biomass in grams.

During the 2022 data collection, two additional traits were captured. An LI-600 (Li-Cor Bioscience, Lincoln, NE, USA) to measure the stomatal conductance and Photosystem II quantum efficiency (ΦPSII) during the solar noon window. Additionally, fresh shoot and root biomass weights were taken prior to drying in 2022. A summary of the measured traits is shown in Table 1.

**Table 1.**
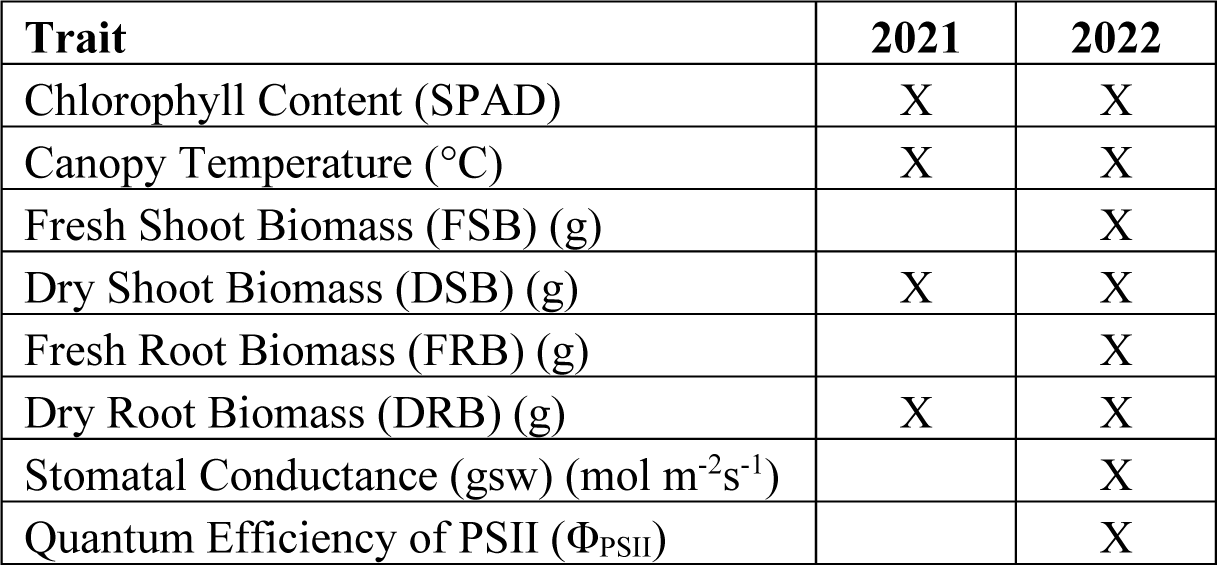
Summary of traits measured in each year of data collection. All traits were measured for both environments (heat and optimal).

### 2.5 Statistical Analysis

Assumptions of analyses of variance (ANOVA) were tested using the Shapiro Wilk test for normality and Barlett’s test using base R functions. Additionally, residuals were visually checked via QQ plots and residual plots for heteroscedasticity to check for heterogeneous variance. Boxcox transformations were used for traits with heteroscedasticity in their residuals. Outliers were removed by calculating studentized residuals for each observation of each trait and excluded from the analysis with values ± 3.4 (Lund, 1975). ANOVA mixed linear model for untransformed phenotypic traits was:

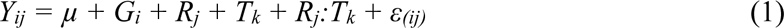

Where *Y_ij_* is the phenotypic value of the *i*^th^ genotype in the *j*^th^ replication, *µ* is the population mean, *G_i_* is the fixed genotypic contribution for the *i*^th^ genotype, *R_j_* is the random replication effect of the *j*^th^ replication, *T_K_* is the fixed treatment effect of the *k*^th^ treatment, *R_j_:T_k_* is the random interaction of the block and treatment effects, and *ε_(ij)_* is the residual error.

Broad sense heritability (*H^2^_Piepho_*) was calculated via the H2cal function in the inti package (Lozano-Isla, 2021) in R and was determined by the equation:

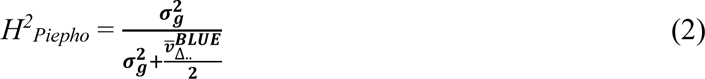

Where 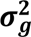 is the genotypic variance and 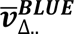 is the average standard error of the genotypic BLUEs.

The best linear unbiased estimates (BLUEs) for each trait was calculated using the H2cal function. A mixed linear model for each condition (heat – 2021, 2022; optimal – 2021, 2022) was fit with the following equation:

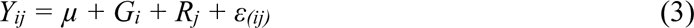

Where *Y_ij_* is the phenotypic value of the *i*^th^ genotype in the *j*^th^ replication, *µ* is the population mean, *G_i_* is the fixed genotypic contribution for the *i*^th^ genotype, *R_j_* is the random replication effect of the *j*^th^ replication, and *ε_(ij)_* is the residual error. Using a completely random model best linear unbiased predictors (BLUPs) for each trait were calculated using (3).

The estimated marginal means (emmeans) of each accession was calculated using the emmeans package in R and fit to the same equation as described in (3). Heat tolerance indices (H_tI_) for all the studied traits were estimated according to the following formula:

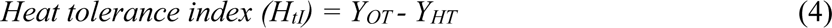

Where *Y_OT_* is the emmean of an accession under the optimal temperature conditions, and *Y_HT_* is the emmean of the same accession e under heat stress temperature conditions. The emmeans of accessions were used to select response to heat (tolerance or susceptibility).

### 2.6 Genome Wide Association Method

Genome-wide association studies (GWAS) were performed using H_tI_ and other traits BLUEs with the SVEN methodology (Li et al., 2023). Of the panel of 450 accessions, 423 had SoySNP50K publicly available (Song et al., 2015), and only these accessions were included in GWAS. The SoySNP50K data has 47K markers across the genome, which were imported into TASSEL and filtered on a minor allele frequency (MAF) of 1%. Genotypic data was imputed and then used to calculate a kinship matrix with Centered IBS in TASSEL. A principal component analysis (PCA) was performed with four components to determine familial or population structure, which was inputted in GWAS. The kinship matrix and PCA from TASSEL were utilized for GWAS in SVEN.

SVEN is a Bayesian method for GWAS based on a hierarchical multi-locus model. It can control for false discovery through prior regularization on the number of important markers (Rairdin et al., 2022; Li et al., 2023). The bravo package (Li et al., 2023) was implemented in R to run SVEN. Given the complexity of abiotic stress traits, we used a larger prior shrinkage lambda of 20. Markers with a marginal inclusion probability (MIP) greater than 0.5 were declared as significant and reported. Candidate gene search was done using the Genome Browser available on Soybase (soybase.org) (Brown et al., 2021) with the Wm82.a2 assembly.

### 2.7 Genomic Prediction (GP)

The genomic best linear unbiased prediction (GBLUP) methodology was utilized for genomic selection (GS) using the rrBLUP package (Endelman, 2011) in R. Phenotypic BLUP data was randomly divided into a training set (80%) and a validation set (20%). The full set of SNP markers from the 423 accessions was used for training, as well as a separate model was trained on a subset of GWAS significant markers that had a p ≤ 0.05 in TASSEL. A ten-fold cross-validation (CV) was performed to avoid inflated estimates of predictive ability. The list of training genotypes was divided into ten equally sized subgroups for the ten-fold CV. Nine of these subgroups were used for training the prediction model, and the remaining one subgroup was used as the testing set. The testing set was used to assess the correlation between the genomic estimated breeding values (GEBVs) of the predicted trait value with the BLUPs of the observed trait value, using the Pearson correlation coefficient. This process was repeated ten times, with each of the subgroups being the testing population once. The process was repeated for 100 cycles by randomly reforming the folds with a different subset of genotypes in each cycle. Across all folds and cycles, the 20% validation set remained unobserved. The correlation accuracy between the GEBVs and BLUPs of the validation population was generated in each cycle. This process was repeated separately for training and predicting in the optimal environment and in the heat environment. The genomic prediction accuracies (GPAs) are reported as the means of the validation (*r_VS_*) and testing (*r_TS_*) correlation coefficients.

Using the developed GP model, the GEBV for each accession in the USDA soybean germplasm collection was calculated for SPAD, DSB, and DRB for both years, and the top 5% (992 accessions) and 10% (1985 accessions) were selected. The top 5% (21 accessions) and 10% (42 accessions) most heat-resistant lines were identified based on their GEBV for all traits for each year. In addition, aiming to understand the diversity between the selected accessions, the country of origin, and the maturity group were verified for the top 5% USDA germplasm. To verify the accessions performance tested in this research with the USDA top accessions, Boxplots were generated to visually compare the 5% and 10% most tolerant accessions identified in the USDA soybean germplasm collection with the 5% and 10% most tolerant accessions identified in the current research.

## 3 RESULTS

### 3.1 Descriptive statistics

The distributions and descriptive statistics for all measured traits are summarized in Figure 1 and Table 2. Phenotypic variation was observed for all traits across a single year, as well as temperature treatment. The effect of genotype was significant for all traits except canopy temperature (p < 0.05). The treatment effect was significant for SPAD, canopy temperature, fresh shoot biomass (FSB), dry shoot biomass (DRB), and stomatal conductance in 2022. The genotype by treatment effect was significant for all traits in 2022 except Φ_PSII_ and canopy temperature (p< 0.05). The interaction effect of genotype and treatment was also significant for DSB and DRB in 2021. Comprehensive ANOVA results are included in Table S2. Heritability estimate was higher in the optimal temperature treatments for most traits in both years (Table 2). The heritability estimates for fresh root and shoot biomass traits were similar across the temperature treatments.

**Figure 1.**
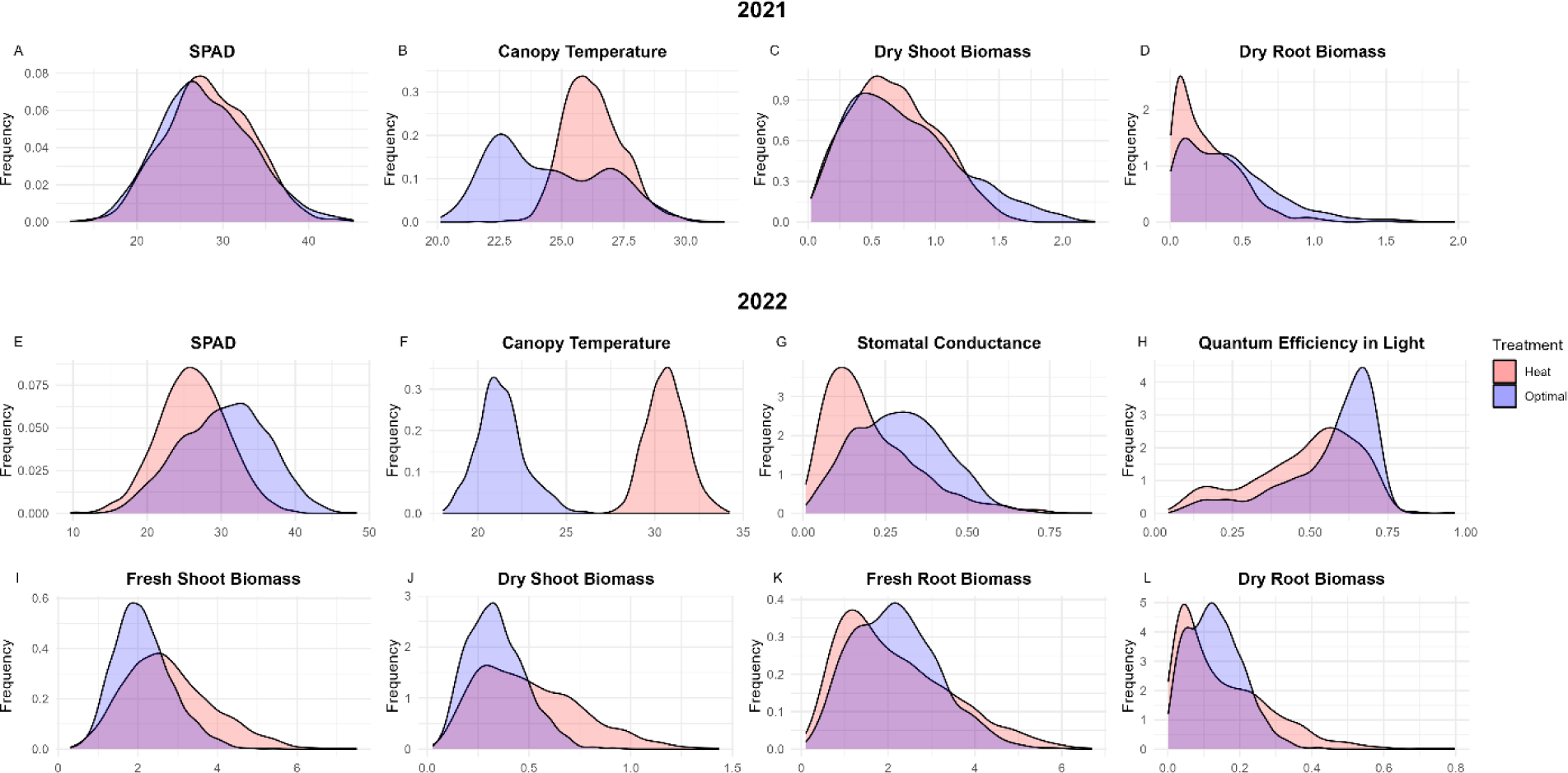
SPAD, Canopy temperature, dry shoot biomass and dry root biomass trait distribution from a greenhouse experiment that included heat and control treatments. Data comes from 450 soybean accessions from the USDA-GRIN mini-core collection. The red is the distribution in heat conditions, and the blue is the distribution in the optimal control conditions. Purple is the overlap of the two distributions.

**Table 2.**
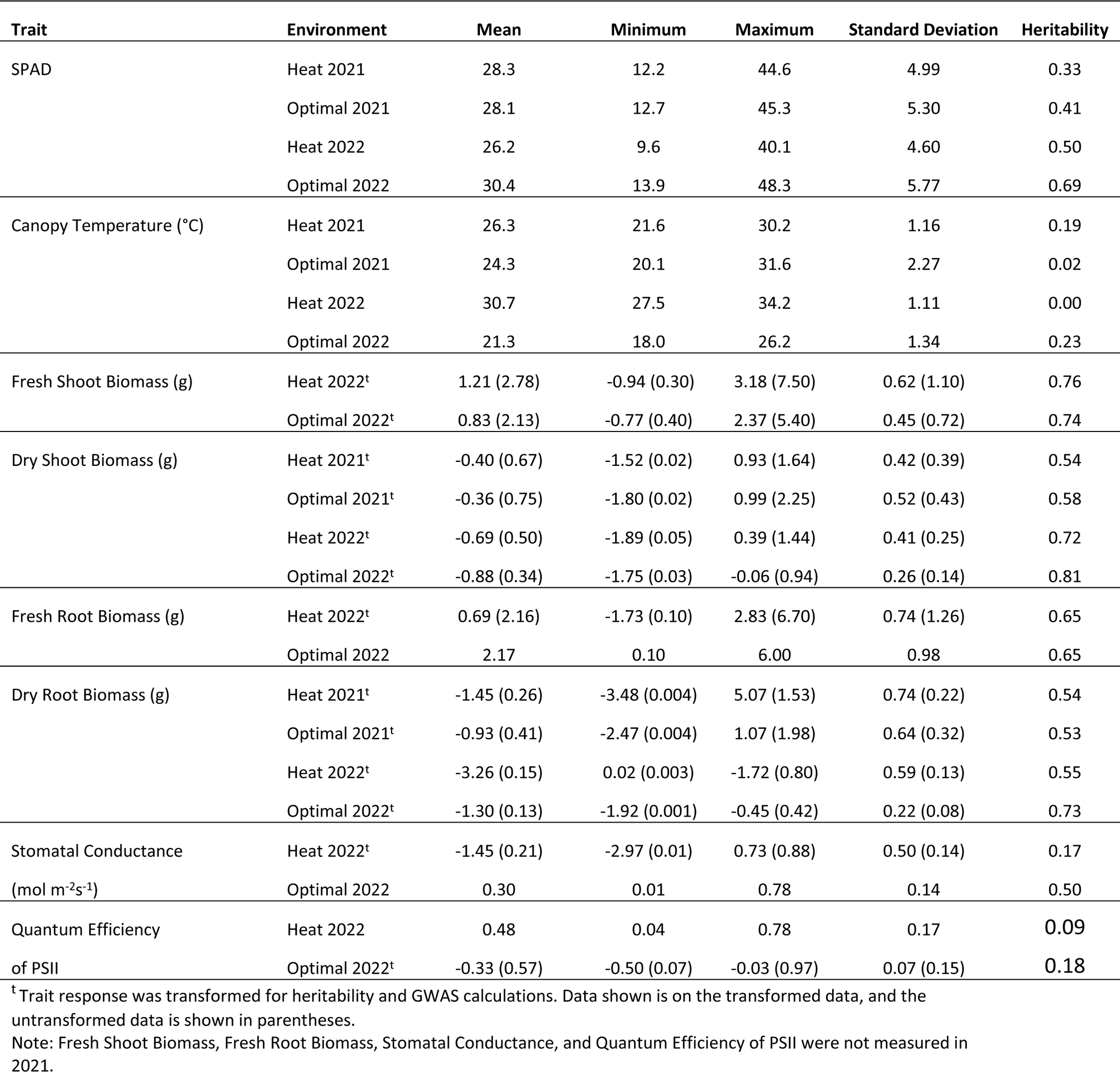
Descriptive statistics and broad sense heritability of all traits measured on the panel of 450 diverse soybean accessions, estimated on four replications in a greenhouse experiment that included heat and control (optimal) treatments.

The chlorophyll content was not different between the heat and optimal conditions in 2021. However, in 2022, chlorophyll content significantly differed between the two temperature conditions, with a lower SPAD value in the heat than in the optimal. Canopy temperature was significantly higher in the heat treatment in both years, with a greater difference observed in 2022 (Table 2). The FSB, DSB and DRB had significantly higher biomass in the heat treatment in 2022, while FRB was not significantly different between the heat and control. The dry biomass traits responded differently in 2021, with significantly lower biomass values in the heat treatment. Both photosynthetic traits, stomatal conductance (gsw) and Φ_PSII_, had significantly lower values in heat compared to the optimal treatment (p < 0.001). For many of the traits, the distribution was wider in the heat treatment as compared to the optimal treatment, indicating a greater diversity of genotypic responses to the higher temperature.

Due to a more comprehensive trait data availability in 2022, we focused on these data to identify five top heat-tolerant soybean accessions. The five heat-tolerant soybean accessions from the USDA-GRIN collection are listed in table 3. These lines were selected based on the HTi values, with a lower HTi better for all traits, excluding canopy temperature. A lower HTi indicates the cultivar’s phenotype was similar or better in the heat treatment as compared to the optimal treatment. The heat tolerant lines are representative of two maturity groups tested in the panel, with the majority from MG II. These lines had phenotypes similar in both temperature treatments or better in the heat for some traits. All of these selected accessions had no significant changes in chlorophyll content, and the majority of the accessions had a significant increase in all biomass traits. The photosynthetic parameter traits had no change between the heat and optimal, except for a higher stomatal conductance rate for PI 437840a.

**Table 3.**
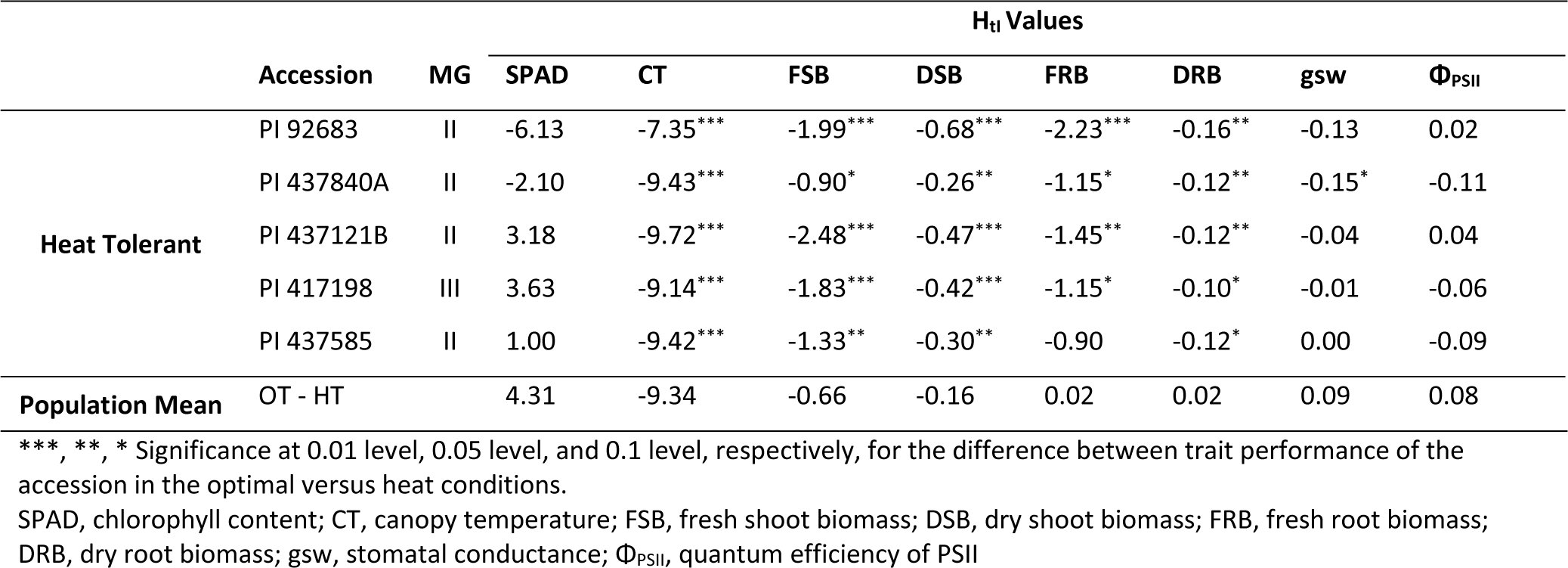
Five PI accessions are classified as heat tolerant. Lower values indicate that the accession performed more similarly or better in heat than in optimal conditions. Higher values indicate a worse performance for a trait in heat conditions. The difference in the population means between the two temperature treatments (OT-HT) is shown for comparison purposes.

For comparison purposes, the difference in population means for the HT and OT treatment are provided to illustrate the trait value differences in these five tolerant accessions relative to the overall 423 accessions. We also investigated five heat susceptible accessions, and noted they generally had lower trait values when grown under heat conditions (Data not presented). The chlorophyll content of these accessions was significantly lower in the heat treatment. These susceptible accessions had lower biomass, and lower photosynthetic parameters with a significant decrease in stomatal conductance for all five susceptible accessions (data not presented).

### 3.2 GENOME WIDE ASSOCIATION STUDY

Association studies detected 37 unique SNPs across different traits and in different temperature treatments (Table 4, Table S4). No significant SNPs were detected in 2021; all further reported GWAS results are from 2022. Table 4 summarizes the distribution of significant SNPs that were identified across traits. The biomass traits had the greatest number of associated SNPs. More SNPs were associated with trait response in the optimal control temperature for most traits. Across all traits, no heat detected SNPs were detected in the optimal treatment for the same trait [NEEDS CLARIFICATION - AKS]. Five of the significant SNPs were associated with more than one trait, with four detected for DRB. A single SNP, *ss715607454*, was detected for HtI traits, with the SNP significant for DSB. This SNP is 3.5 kbp away from *Glyma.10G219100* and 9.1 kbp away from *Glyma.10G219200*. Both genes are described as encoding for laccase proteins, which are glycoproteins involved in the production of lignin for plant growth and have been found to be involved with abiotic stress response in several species (Bai et al., 2023). In addition to these nearby candidate genes, the gene *Gylma.10G219600* is 37.4 kbp downstream and encodes for an NAC transcription factor, *GmNAC074*, a class of proteins previously reported to be involved in response to abiotic stress in soybean (Tran et al., 2009; Melo et al., 2018). Further upstream, at 135.8 kbp away, *Glyma.10G217900* has been reported to be upregulated in soybeans grown in heat-stress conditions (Wang et al., 2018).

**Table 4.**
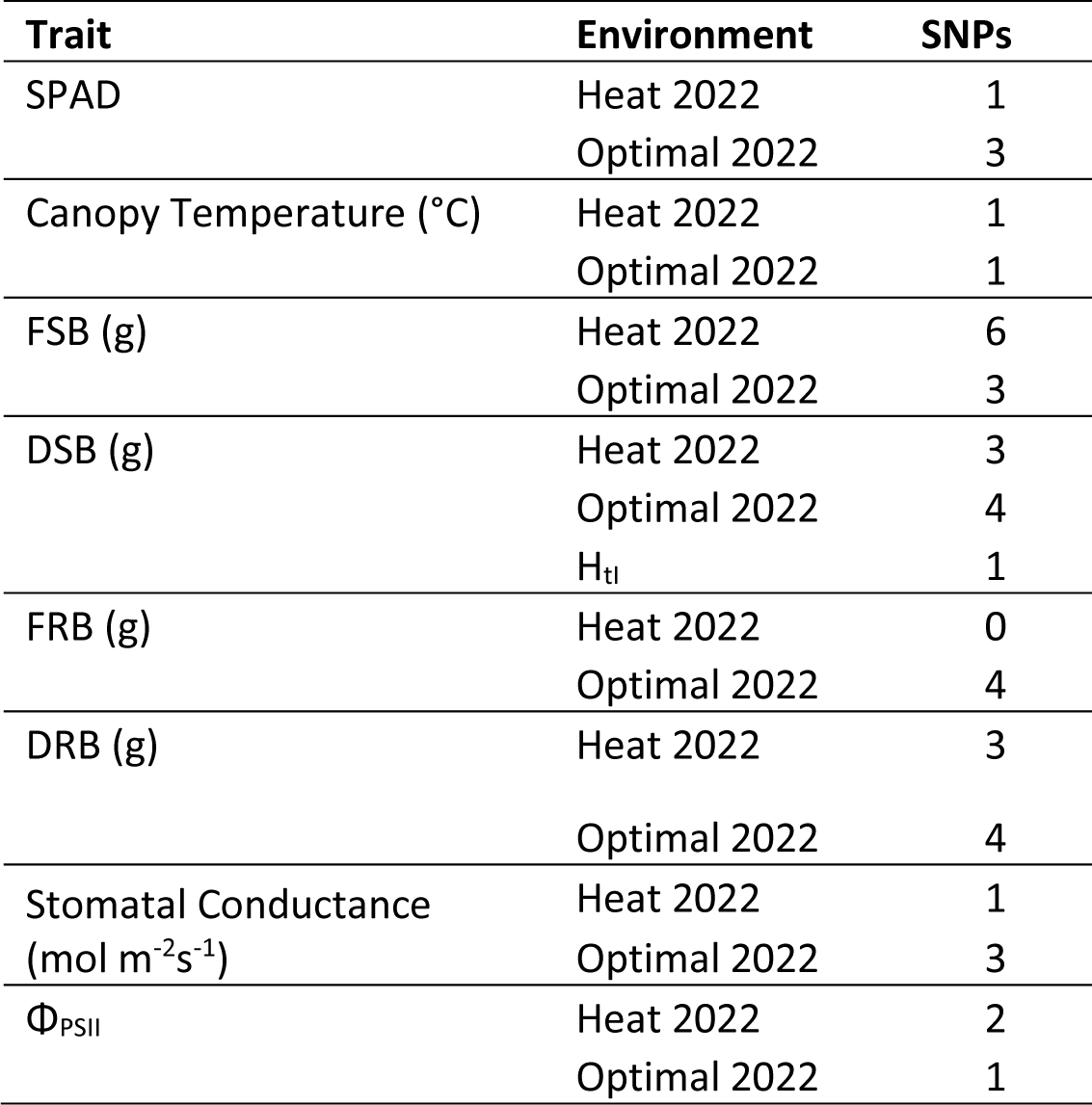
Summary of the significant SNPs associated with each trait and environment, using the SVEN GWAS methodology.

Table 5 lists a subset of potential candidate genes from the SNPs we detected. The SNP *ss715588916* was detected in the heat treatment and associated with FSB and DRB. This SNP is 61.9 kbp away from *Glyma.04G242300*, which has been previously reported to be downregulated in soybean due to heat (Wang et al., 2018) and encodes for a plantacyanin protein. The SNP *ss715604755* was detected in the optimal conditions for the two fresh biomass traits. This SNP is within *Glyma.09G249100*, a glutamine-fructose-6-phopshate transaminase (GFAT). In optimal conditions, two SNPs, *ss715613671* and *ss715612610*, were associated with FSB and DRB. The latter SNP is 5.4 kbp from the candidate gene *Glyma.12G197600*, a xyloglucan endotransglucosylase/hydrolase (XTH) protein. A single SNP was detected across temperature treatments, although in different traits. This SNP, *ss715627447*, was found to be significant for FSB in the heat treatment and optimal for DRB. It is 49.8 kbp upstream from *Glyma.17G228800*, which has been reported to be upregulated in soybean exposed to heat stress (Wang et al., 2018a) and is a glycine decarboxylase P-protein.

**Table 5.**
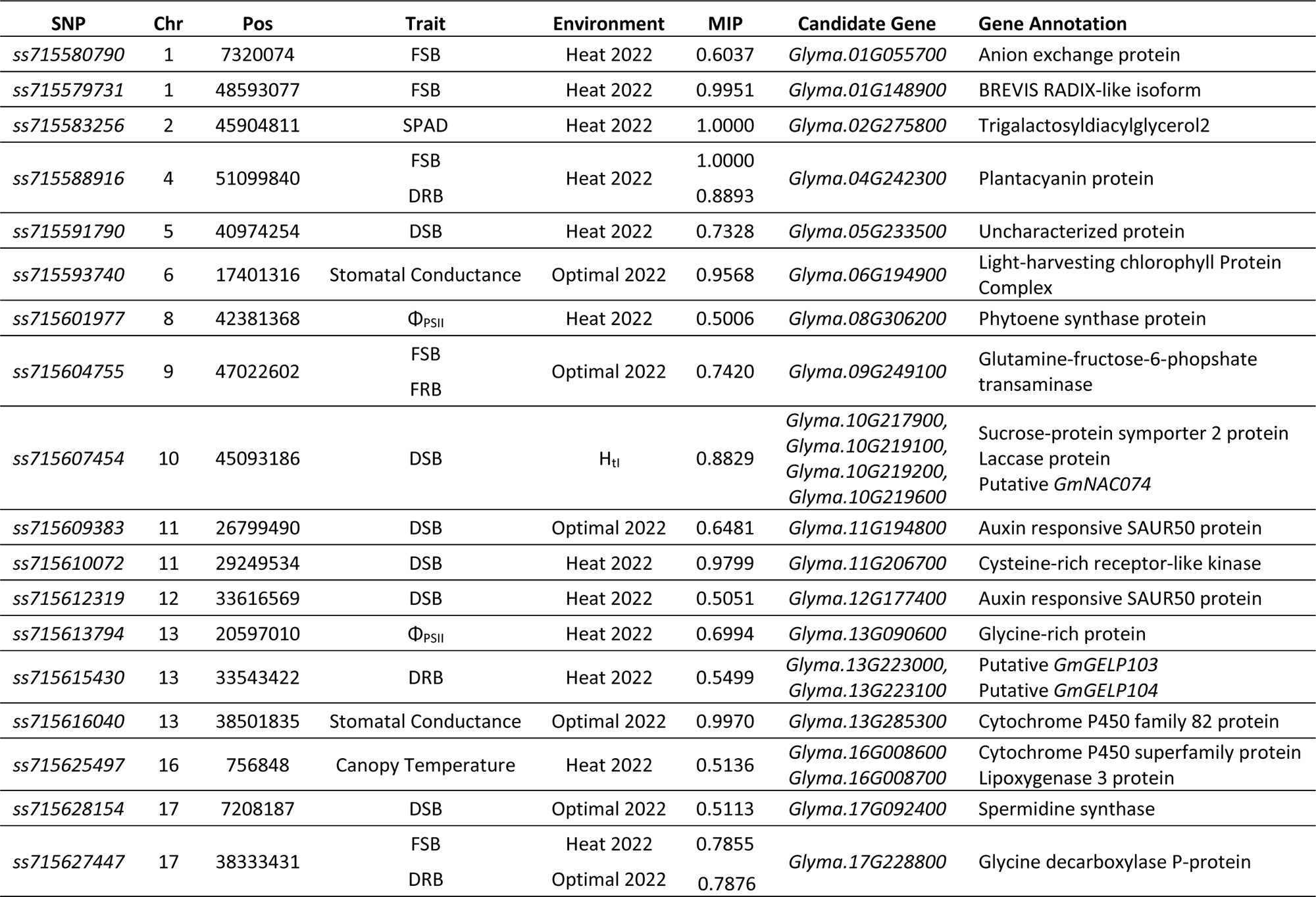
Description of a subset of significant marker-trait associations from the SVEN model, and the candidate genes associated with them inferred from the soybean genome Wm82.a2. The MIP and environment that the SNP was detected in is included.

The candidate gene *Glyma.01G148800* is 9.9 kbp away from *ss715579731*, a SNP reported in association with FSB in heat. This gene encodes for an uncharacterized protein and has been previously found to be downregulated in soybeans exposed to heat stress (Wang et al., 2018). This SNP is also within the candidate gene *Glyma.01G148900*, which is a Brevis radix-like (BRX) protein. Another potential candidate gene on Chromosome 2 is *Glyma.02G238300*, a mitochondrial FE/S cluster exporter protein previously reported as significant for total chlorophyll content (Dhanapal et al., 2016). This gene is 157.4 kbp away from *ss715583256*, which was associated with SPAD in heat. The SNP is also 25.6 kbp away from an additional potential candidate gene, *Glyma.02G275800*, a trigalactosyldiacylglycerol2 (TGD2) protein. The gene *Glyma.06G194900* is 17.2 kbp away from a SNP, *ss715593740*, associated with stomatal conductance in the optimal treatment. This candidate gene is a light-harvesting chlorophyll-protein complex. Another SNP associated with stomatal conductance, *ss715616040*, is downstream of *Glyma.13G285300*, which is a cytochrome P450 (CYP450) family protein. The SNP *ss715601977*, which is associated with ΦPSII in heat, is 40.9 kbp away from the candidate gene *Glyma.08G306200*, which is a phytoene synthase protein. Another SNP associated with ΦPSII in heat, *ss715613794*, is 42.3 kbp away from the candidate gene *Glyma.13G090600*, which encodes for a glycine-rich protein (GRP) and has been found to be upregulated in heat (Wang et al., 2018).

### 3.3 Genomic Prediction

Genomic prediction ability (GPA) was calculated against the testing and unseen validation sets. Training was performed on 2021, 2022, and combined year data when available. Canopy temperature was dropped from genomic prediction analysis due to the non-significant genotypic effect. A summary of the GPAs is shown in Table 6. The GPA for most traits under heat conditions decreased when tested against the unseen validation set compared to the testing set. In three cases (2021 DSB, 2021 DRB, 2022 ΦPSII), the GPA against the validation set was higher than against the testing set. In traits with two years of data, there was a trend of higher GPA in the 2022 data, which follows a trend of higher heritability in the 2022 data. For SPAD, the highest GPA came from combining data from the two years, while this combined data resulted in a lower GPA for DSB and DRB. The same trend of higher GPA for the test set was true when trained on the traits grown in the control. Only the 2022 SPAD and the 2022 ΦPSII had higher GPAs when tested against the validation set. For all traits except 2022 ΦPSII, the GPA for the test set was higher when trained and tested on data grow in the control compared to the heat, although this increase was non-significant in most cases. The GPA for the validation set was higher in the control than heat, except in 2022 FSB, 2021 DSB, and 2022 ΦPSII. This trend of higher GPA in the control follows with the higher heritabilities found in the control set.

**Table 6.** The genomic prediction accuracies of each trait-environment combination using two different marker densities (1) the full SNP panel and (2) the SNPs with a p ≤ 0.05 via TASSEL.

| Trait | Year | Full Panel SNPs |  |  |  | Marker-Trait Associated SNPs |  |  |  |  |  |
| --- | --- | --- | --- | --- | --- | --- | --- | --- | --- | --- | --- |
|  |  | Heat |  | Optimal |  | Heat |  | Optimal |  | Heat<sup>§</sup> |  |
| | | $r_{TS}$ | $r_{VS}$ | $r_{TS}$ | $r_{VS}$ | $r_{TS}$ | $r_{VS}$ | $r_{TS}$ | $r_{VS}$ | $r_{TS}$ | $r_{VS}$ |
| SPAD | 2021 | 0.31 | 0.12 | 0.34 | 0.26 | 0.33 | 0.15 | 0.22 | 0.26 | 0.31 | 0.21 |
|  | 2022 | 0.32 | 0.13 | 0.44 | 0.50 | 0.28 | 0.12 | 0.39 | 0.50 | 0.31 | 0.18 |
|  | Combined | 0.39 | 0.14 | 0.53 | 0.39 | 0.38 | 0.16 | 0.55 | 0.37 | 0.36 | 0.22 |
| Fresh Shoot Biomass | 2022 | 0.61 | 0.53 | 0.51 | 0.51 | 0.55 | 0.47 | 0.43 | 0.48 | 0.54 | 0.42 |
| Dry Shoot Biomass | 2021 | 0.35 | 0.40 | 0.47 | 0.37 | 0.33 | 0.40 | 0.42 | 0.36 | 0.33 | 0.31 |
|  | 2022 | 0.58 | 0.50 | 0.59 | 0.54 | 0.51 | 0.50 | 0.55 | 0.53 | 0.53 | 0.42 |
|  | Combined | 0.54 | 0.44 | 0.54 | 0.52 | 0.51 | 0.40 | 0.52 | 0.47 | 0.51 | 0.39 |
| Fresh Root Biomass | 2022 | 0.49 | 0.41 | 0.54 | 0.51 | 0.45 | 0.37 | 0.50 | 0.48 | 0.46 | 0.37 |
| Dry Root Biomass | 2021 | 0.34 | 0.38 | 0.42 | 0.36 | 0.30 | 0.34 | 0.40 | 0.37 | 0.32 | 0.39 |
|  | 2022 | 0.52 | 0.43 | 0.59 | 0.56 | 0.47 | 0.38 | 0.58 | 0.54 | 0.48 | 0.39 |
|  | Combined | 0.49 | 0.41 | 0.54 | 0.52 | 0.45 | 0.36 | 0.53 | 0.48 | 0.47 | 0.39 |
| Stomatal Conductance | 2022 | 0.21 | 0.10 | 0.45 | 0.31 | 0.17 | 0.15 | 0.31 | 0.29 | 0.16 | 0.09 |
| Quantum Efficiency of PSII | 2022 | 0.15 | 0.27 | 0.11 | 0.22 | 0.12 | 0.20 | -0.02 | 0.18 | 0.16 | 0.32 |

<sup>§</sup>SNPs were selected from the optimal environment TASSEL results and were used for training a genomic prediction model for prediction in a heat-stress environment.

The number of marker-trait associated SNPs used for different trait and year combinations varied between 933 and 2289. The mean accuracy of the predictions based on trait associated SNPs obtained through GWAS was lower than the mean accuracy when using the entirety of the 50K SNP panel (Table 6). 2021 Heat SPAD was the only trait with a higher prediction accuracy for both the testing and validation sets when the trait associated SNPs were used for training. Interestingly, when the SNPs detected in the control conditions were used to train to predict traits grown in heat, the prediction accuracy was not different from the heat selected SNPs.

Based on data availability across years and GPAs, three traits were selected for use in germplasm projection: SPAD, DSB, and DRB. The average trait performance of the top 5% and 10% accessions in the USDA collection showed a higher average across the accessions evaluated in this study (Figure 2), indicating that it is possible to find other heat tolerant accessions in the USDA germplasm collection. The top 5% and top 10% of heat tolerant accessions in the panel we screened overlapped with the top 5% accessions when projected to the entirety of the USDA collection. For the traits, there was an overlap of 100%, 80%, and 90% for SPAD, DSB, and DRB, respectively (Figure 2). The country of origin of the top 5% heat tolerant accession from the germplasm collection varied, with China, United States, Japan, South Korea, and Russia as the five countries contributing the most (Figure S2). The maturity groups of the top 5% heat tolerant accessions was mainly in the early maturity groups 0-II (Figure S3).

**Figure 2.**
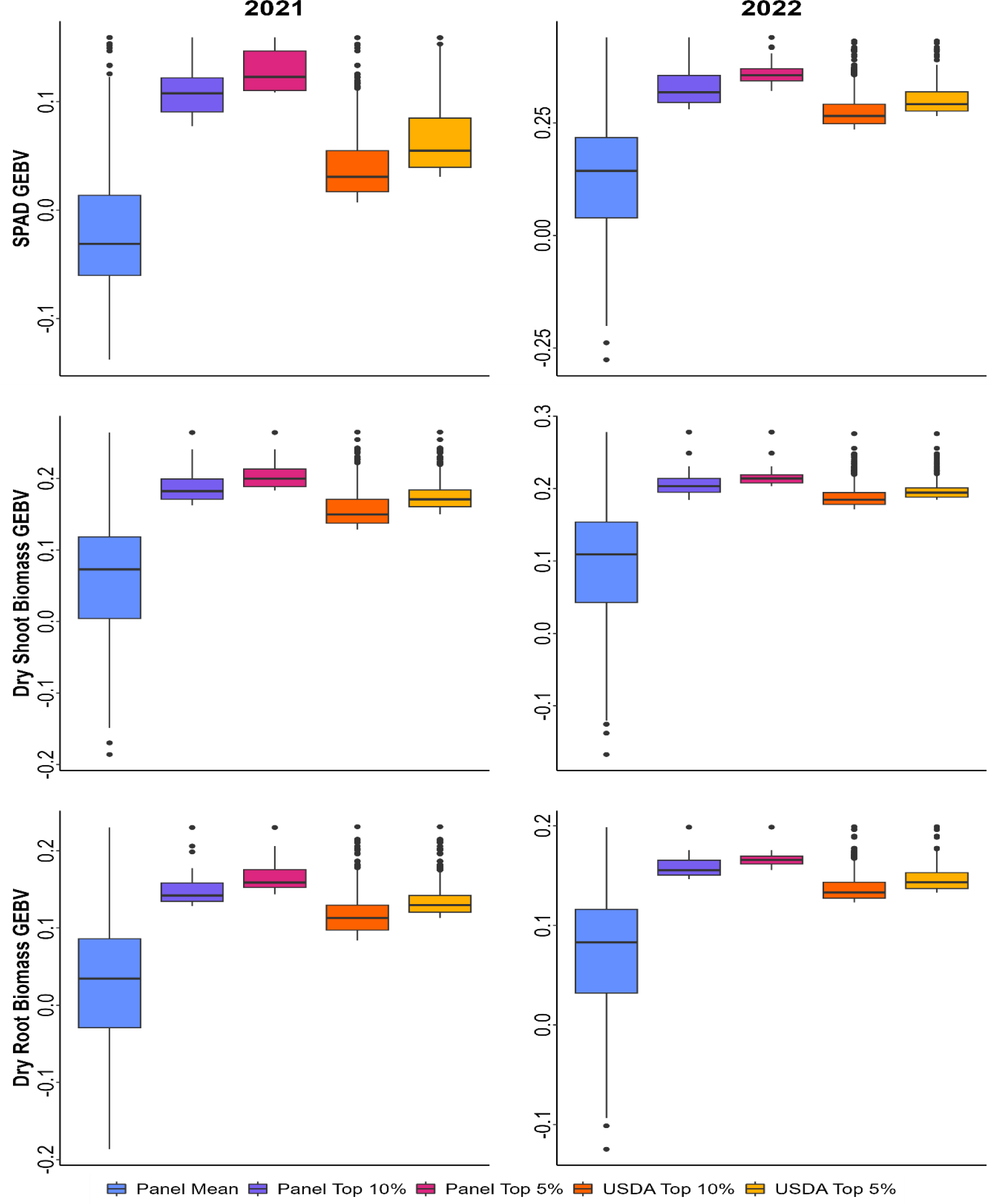
Genetic estimated breeding value for the 423 accessions included in this study (Panel Mean), 10% highest for the 423 accession evaluated (Panel Top 10%), 5% highest for the 423 accession evaluated (Panel Top 5%), 10% highest for the entire USDA germplasm collection (USDA Top 10%), 5% highest for the entire USDA germplasm collection (USDA Top 5%) for SPAD, DSB, and DRB for two years (2021 and 2022).

## 4. DISCUSSION

Our study leveraged the USDA-GRIN soybean mini-core collection to obtain detailed trait data in response to heat stress. In this first step, we focused on the early growth stages stress response to aid in the identification of useful access for breeding applications, generate genetic insights, and enhance phenomics-assisted selection (Singh et al., 2021). Our study revealed that heat stress adversely affected physiological traits such as chlorophyll content (SPAD), stomatal conductance, and Quantum Efficiency of PSII, similar to what was reported with previous studies (Poudel et al., 2023). Additionally, canopy temperatures increased under heat stress consistent with previous report (Poudel et al., 2023). The change in biomass to heat was inconsistent over two years; 2021 saw reduced biomass under heat stress, while 2022 exhibited increased biomass. In a separate near-field abiotic stress screening (N-FAST) of eight soybean varieties, we found that there was no significant change in shoot biomass (15 g plant^−1^ in control and 14.3 g plant^−1^ in heat) and a significant increase of 49.6% in root biomass (8.2 g plant ^−1^ in control and 12.3 g plant^−1^ in heat) due to heat stress. The increase in biomass under heat can possibly be due to accelerated growth stages and internode elongation, which has been previously reported in tomato (Zhou et al., 2017). We noted seven accessions had initiated flowering (R1 growth stage) within three weeks of plants, which is dramatically different from the normal expectation of planting to R1 duration of about 6-8 weeks. Generally, heritability was lower under heat stress, a trend also observed in other crops under marginal conditions (Al-Yassin et al., 2005; Kumar et al., 2007; Longmei et al., 2021).

Understanding the genetic architecture of physiological traits evaluated under heat stress helps identify markers and develop genomic selection pipelines to assist in the development of heat tolerant soybean. To date, very limited studies have reported GWAS in soybean that reported on tolerance to heat stress. To explore heat stresses’ complex genetic architecture using GWAS involving 423 diverse soybean accessions, we measured eight traits related to growth and physiological parameters in an optimal control temperature and in a heat stress temperature. We identified 37 unique SNPs that were associated with different traits and distributed across 17 of the 20 soybean chromosomes. SNPs associated with traits in the optimal treatment made up 20 of the detected SNPs, with 15 of the additional SNPs detected in heat, one SNP detected across temperature treatments, and the remaining one SNP detected for an H_tI_. This lack of overlap indicates a genetic divergence in soybean growth and development under optimal conditions and in stress conditions, which has been previously reported in corn (Yuan et al., 2019; Longmei et al., 2021).

In the current study, several putative candidate genes were identified for the physiological traits in heat stress and without stress. The SNP detected for the H_tI_ DSB, *ss715607454*, is of particular interest due to the four potential candidate genes in the region. These genes includes two laccase proteins, a NAC transcription factor, and a sucrose-protein symporter. All of these proteins have been found to be involved in the response to abiotic stress in soybean and other plant species (Tran et al., 2009; Melo et al., 2018; Soltani et al., 2019; Bai et al., 2023)). The SNP associated with Φ_PSII_ in heat, *ss715601977* is near a phytoene synthase protein gene. Phytoene synthase is essential for modulating the biosynthesis of the carotenoid pigment (Zhou et al., 2022), which are a light-harvesting pigment used in the photosystem complexes and are also essential for protection against reactive oxygen species produced in the chloroplast due to stress (Young, 1991).

The biomass traits have several candidate genes of particular interest. The SNP *ss715579731*, is within a gene that encodes for a BRX protein, which is a class of proteins that have been found to promote shoot and root growth in Arabidopsis (Beuchat et al., 2010) and rice (Li et al., 2019). Two SNPs associated with DSB in heat have candidate genes of interest. The first SNP, *ss715591790*, is near the gene *Glyma.05G233500*, which encodes for an unknown protein that was downregulated in soybeans exposed to heat stress (Wang et al., 2018). The second SNP, *ss715610072*, is near the candidate gene *Glyma.11G206700*. This gene is a cysteine-rich receptor-like kinase (CRK), which is a family of receptor-like kinases (RLKS) that are important in plant immunity, response to abiotic stress, and regulating growth and development (Zhang et al., 2023). CRKs are highly evolutionarily conserved across vascular plants and have been found to modulate the tradeoff between growth and stress response in many species (Zhang et al., 2023). The SNP on chromosome 13 associated with DRB in heat has two promising candidate genes nearby, *Glyma.13G223000* and *Glyma.13G223100*. These two genes are the *GmGELP103* and *GmGELP104* genes (Su et al., 2020), which are GDSL-type esterase/lipase proteins. The *GDSL* gene family has been found to be involved in growth and development as well as response to stress in many plant species, including the identification of *GmGELP* genes as candidates for salinity and drought tolerance in soybean (Su et al., 2020). The numerous significant SNP points to the complexity of heat tolerance response in soybean. In other crops, namely, common bean (López-Hernández and Cortés, 2019), peanut (Sharma et al., 2023), maize (Longmei et al., 2021; Seetharam et al., 2021), and sorghum (Chen et al., 2017), a large number of significant SNP were reported that control heat tolerance. While these multitude of SNP and associated genes are useful to decipher the genetic control of heat tolerance, they have limitation in immediate application in heat tolerant variety development due to their large number and smaller effects. Therefore, there is a need to explore genomic prediction methods to investigate their usefulness in genomic selection.

Genomic prediction is a powerful tool in plant breeding (Crossa et al., 2017), with the ability to estimate the breeding value of accessions without having to use the resources to phenotype in greenhouses or fields. Genomic prediction has been found in soybean to have a higher predictive ability than phenomic selection alone (Howard and Jarquin, 2019). We evaluated the effectiveness of genomic selection for soybean biomass and physiological processes in heat versus optimal temperature conditions. We were also interested in using GP to project to the USDA GRIN collection that has been previously shown to be effective in soybean (de Azevedo Peixoto et al., 2017) and maize (Yu et al., 2016). In this study, we observed that GPA was higher for genomic selection applied at the optimal temperature compared to the heat treatment. The GPA is largely dependent on several factors, including linkage disequilibrium (LD), number of markers, heritability, and population size. Given that the same population with the same number of markers was used for training, the differences in GPA can largely be attributed to the differences in trait heritability, which were lower in heat treatment and consistent with previous studies (Ud-Din et al., 1992; Kumar et al., 2007). The GPA for the four biomass traits remained comparable across the heat and optimal treatments, indicating that genomic selection for biomass is feasible in heat conditions.

Given the moderate predictive ability of the biomass traits in heat, we projected the GP model to estimate GEBVs for the entire USDA soybean germplasm collection. Given that the soybean collection has more than 20,000 accessions available, it is not feasible to phenotype all accessions for heat stress tolerance. By projecting the GP model, GEBVs can be estimated, which can be used by soybean breeders to select genotypes to be used in crossing for heat stress tolerant varieties enabling them to utilize accessions appropriate to their maturity group. The germplasm projection trended towards lower maturity groups for a large proportion of the heat tolerant accession. This could be due to the model being trained primarily on these maturity groups or could indicate a differential response to heat stress between early and later maturity groups. The efforts to utilize germplasm collection need to be increased as they hold tremendous value in improving traits to combat the adverse effects of climate change.

The results from our study can benefit breeding efforts aimed at early season heat response, as the accessions we report are targeted to the early growth stage. Additional studies are needed to measure heat stress response in critical growth stages, such as flowering, pod and seed initiation. Large-scale screening for heat stress remains a bottleneck due to the difficulty of phenotype accessions in the field with the option to compare them to optimal temperature treatment. Additional difficulty arises as the areas projected to experience temperature increase in the future do not provide fields with natural heat stress, while areas with higher current temperatures are in dissimilar maturity zone and soybean accessions will experience a confounding photoperiod response. Solutions for screening for soybean heat stress tolerance could include utilization of large indoor spaces, construction of artificial heat sources in specialized fields, and development of phenotyping traits to be used in indirect selection of heat tolerance. Traits identified in this study should be explored further in near field conditions and developed so that HTP methods can be applied for screening heat tolerance in large soybean breeding populations.

## CONCLUSION

Our study found that there was significant phenotypic and genotypic variability in soybean response to heat stress in the early season. We also identified phenotypes that could be adapted to field-scale HTP methods, which would allow soybean breeders in the future to screen large amounts of breeding materials for heat stress tolerance in the field. The existence of this natural genetic diversity and a preliminary genetic architecture for traits measured in a heat stress environment indicate that these traits can be improved through breeding strategies. Newer breeding strategies, such as genomic prediction, are valuable tools to overcome the difficulty of phenotyping large numbers of accessions for heat stress tolerance in a field setting. These techniques should be validated in the field in future studies to better understand the reaction of soybeans in a heat-stress setting in otherwise natural field conditions.

## Abbreviations

BLUEs: Best linear unbiased estimates
BLUPs: best linear unbiased predictor
DRB: dry root biomass
DSB: dry shoot biomass
emmeans: estimated marginal mean
FRB: fresh root biomass
FSB: fresh shoot biomass
GP: genomic prediction
GWAS: genome-wide association study
H_tI_: heat tolerance index
MIP: marginal inclusion probability
SNP: single nucleotide polymorphism
SVEN: Selection of Variables with Embedded screening using Bayesian methods
TASSEL: Trait analysis by ASSociation, Evolution, and Linkage

## DATA AVAILBILITY STATEMENT

Data is available upon request.

## AUTHOR CONTRIBUTIONS

LV and AKS conceptualized and designed the experiment. LV performed data collection with the assistance of lab members, performed the data analysis, and wrote the first draft manuscript with feedback from AKS. LAP performed the genomic projection analysis and provided feedback on the draft manuscript.

## ACKNOWLEDGMENTS

We thank the undergraduate and graduate students for their assistance during the data collection. We express our gratitude to Jennifer Hicks and Antonella Ferela for their assistance in procuring materials and organizing assistance for this project. We thank Aaron Brand and Peter Lawlor for ensuring the greenhouses were running in the appropriate settings. We appreciate support from Dr. Phillip Dixon and Ms. Laritza Lorenzo for their guidance and feedback on the statistical analysis, and Dr. Jacqueline Campbell for assistance in the search of candidate genes.

## SUPPLEMENTAL MATERIAL

Supplemental materials include five tables and two figures.

Figure S1. Distribution of the country of origin for the top 5% most tolerant soybean accessions from the genomic projection.

Figure S2. Distribution of the maturity group for the top 5% most tolerant soybean accession from the genomic projection.

Table S1. The complete list of 450 soybean accessions screened in the study, including the maturity group and country of origin.

Table S2. Calendar of planting and phenotyping dates.

Table S3. Complete ANOVA results of the fixed effects for the mixed linear models of each phenotypic trait in each year.

Table S4. Complete listing of significant SNPs detected by SVEN, and their position in the genome.

Table S5. Comparison of the SVEN methodology with the TASSEL and FarmCPU methodologies.

## REFERENCES

Abou-Elwafa, S.F., and T. Shehzad. 2021. Genetic diversity, GWAS and prediction for drought and terminal heat stress tolerance in bread wheat (Triticum aestivum L.). Genet. Resour. Crop Evol. 68(2): 711–728.

Almeselmani, M., P.S. Deshmukh, and V. Chinnusamy. 2012. Effects of prolonged high temperature stress on respiration, photosynthesis and gene expression in wheat (Triticum aestivum L.) varieties differing in their thermotolerance. Plant stress 6(1): 25–32.

Alsajri, F.A., B. Singh, C. Wijewardana, J.T. Irby, W. Gao, et al. 2019. Evaluating Soybean Cultivars for Low- and High-Temperature Tolerance During the Seedling Growth Stage. Agronomy 9(1): 13.

Alsajri, F.A., C. Wijewardana, J.T. Irby, N. Bellaloui, L.J. Krutz, et al. 2020. Developing functional relationships between temperature and soybean yield and seed quality. Agron. J. 112(1): 194–204.

Al-Yassin, A., S. Grando, O. Kafawin, A. Tell, and S. Ceccarelli. 2005. Heritability estimates in contrasting environments as influenced by the adaptation level of barley germ plasm. Ann. Appl. Biol. 147(3): 235–244.

Assefa, T., J. Zhang, R.V. Chowda-Reddy, A.N. Moran Lauter, A. Singh, et al. 2020. Deconstructing the genetic architecture of iron deficiency chlorosis in soybean using genome-wide approaches. BMC Plant Biol. 20(1): 42.

de Azevedo Peixoto, L., T.C. Moellers, J. Zhang, A.J. Lorenz, L.L. Bhering, et al. 2017. Leveraging genomic prediction to scan germplasm collection for crop improvement. PLoS One 12(6): e0179191.

Bai, Y., S. Ali, S. Liu, J. Zhou, and Y. Tang. 2023. Characterization of plant laccase genes and their functions. Gene 852: 147060.

Balla, K., I. Karsai, P. Bónis, T. Kiss, Z. Berki, et al. 2019. Heat stress responses in a large set of winter wheat cultivars (Triticum aestivum L.) depend on the timing and duration of stress. PLoS One 14(9): e0222639.

Beuchat, J., E. Scacchi, D. Tarkowska, L. Ragni, M. Strnad, et al. 2010. BRX promotes Arabidopsis shoot growth. New Phytol. 188(1): 23–29.

Bita, C.E., and T. Gerats. 2013. Plant tolerance to high temperature in a changing environment: scientific fundamentals and production of heat stress-tolerant crops. Front. Plant Sci. 4: 273.

Brown, A.V., S.I. Conners, W. Huang, A.P. Wilkey, D. Grant, et al. 2021. A new decade and new data at SoyBase, the USDA-ARS soybean genetics and genomics database. Nucleic Acids Res. 49(D1): D1496–D1501.

Chen, J., R. Chopra, C. Hayes, G. Morris, S. Marla, et al. 2017. Genome-wide association study of developing leaves’ heat tolerance during vegetative growth stages in a sorghum association panel. Plant Genome 10(2). doi: 10.3835/plantgenome2016.09.0091.

Cohen, I., S.I. Zandalinas, C. Huck, F.B. Fritschi, and R. Mittler. 2021. Meta-analysis of drought and heat stress combination impact on crop yield and yield components. Physiol. Plant. 171(1): 66–76.

Crossa, J., P. Pérez-Rodríguez, J. Cuevas, O. Montesinos-López, D. Jarquín, et al. 2017. Genomic Selection in Plant Breeding: Methods, Models, and Perspectives. Trends Plant Sci. 22(11): 961–975.

Deshmukh, R., H. Sonah, G. Patil, W. Chen, S. Prince, et al. 2014. Integrating omic approaches for abiotic stress tolerance in soybean. Front. Plant Sci. 5: 244.

Dhanapal, A.P., J.D. Ray, S.K. Singh, V. Hoyos-Villegas, J.R. Smith, et al. 2016. Genome-wide association mapping of soybean chlorophyll traits based on canopy spectral reflectance and leaf extracts. BMC Plant Biol. 16(1): 174.

Dobbels, A.A., and A.J. Lorenz. 2019. Correction to: Soybean iron deficiency chlorosis high throughput phenotyping using an unmanned aircraft system. Plant Methods 15: 113.

Endelman, J.B. 2011. Ridge regression and other kernels for genomic selection with R package rrBLUP. Plant Genome 4(3): 250–255.

Gray, S.B., O. Dermody, S.P. Klein, A.M. Locke, J.M. McGrath, et al. 2016. Intensifying drought eliminates the expected benefits of elevated carbon dioxide for soybean. Nat Plants 2(9): 16132.

Hatfield, J.L., K.J. Boote, B.A. Kimball, L.H. Ziska, R.C. Izaurralde, et al. 2011. Climate impacts on agriculture: Implications for crop production. Agron. J. 103(2): 351–370.

Hatton, N., A. Sharda, W. Schapaugh, and D. van der Merwe. 2020. Remote thermal infrared imaging for rapid screening of sudden death syndrome in soybean. Comput. Electron. Agric. 178(105738): 105738.

Hesketh, J.D., D.L. Myhre, and C.R. Willey. 1973. Temperature control of time intervals between vegetative and reproductive events in soybeans^1^. Crop Sci. 13(2): 250–254.

Howard, R., and D. Jarquin. 2019. Genomic Prediction Using Canopy Coverage Image and Genotypic Information in Soybean via a Hybrid Model. Evol. Bioinform. Online 15: 1176934319840026.

IPCC. 2023. Food, Fibre and Other Ecosystem Products. Climate Change 2022 – Impacts, Adaptation and Vulnerability: Working Group II Contribution to the Sixth Assessment Report of the Intergovernmental Panel on Climate Change. Cambridge University Press. p. 713–906

Jagadish, S.V.K., D.A. Way, and T.D. Sharkey. 2021. Plant heat stress: Concepts directing future research. Plant Cell Environ. 44(7): 1992–2005.

Jin, Z., E.A. Ainsworth, A.D.B. Leakey, and D.B. Lobell. 2018. Increasing drought and diminishing benefits of elevated carbon dioxide for soybean yields across the US Midwest. Glob. Chang. Biol. 24(2): e522–e533.

Jumrani, K., and V.S. Bhatia. 2019. Interactive effect of temperature and water stress on physiological and biochemical processes in soybean. Physiol. Mol. Biol. Plants 25(3): 667–681.

Kaler, A.S., J.D. Ray, W.T. Schapaugh, C.A. King, and L.C. Purcell. 2017. Genome-wide association mapping of canopy wilting in diverse soybean genotypes. Theor. Appl. Genet. 130(10): 2203–2217.

Kidokoro, S., K. Watanabe, T. Ohori, T. Moriwaki, K. Maruyama, et al. 2015. Soybean DREB1/CBF-type transcription factors function in heat and drought as well as cold stress-responsive gene expression. Plant J. 81(3): 505–518.

Kumar, R., R. Venuprasad, and G.N. Atlin. 2007. Genetic analysis of rainfed lowland rice drought tolerance under naturally-occurring stress in eastern India: Heritability and QTL effects. Field Crops Res. 103(1): 42–52.

Leff, B., N. Ramankutty, and J.A. Foley. 2004. Geographic distribution of major crops across the world. Global Biogeochem. Cycles 18(1). doi: 10.1029/2003gb002108.

Li, D., S. Dutta, and V. Roy. 2023. Model Based Screening Embedded Bayesian Variable Selection for Ultra-high Dimensional Settings. J. Comput. Graph. Stat. 32(1): 61–73.

Li, Z., Y. Liang, Y. Yuan, L. Wang, X. Meng, et al. 2019. OsBRXL4 Regulates Shoot Gravitropism and Rice Tiller Angle through Affecting LAZY1 Nuclear Localization. Mol. Plant 12(8): 1143–1156.

Li, P.-S., T.-F. Yu, G.-H. He, M. Chen, Y.-B. Zhou, et al. 2014. Genome-wide analysis of the Hsf family in soybean and functional identification of GmHsf-34 involvement in drought and heat stresses. BMC Genomics 15(1): 1009.

Longmei, N., G.K. Gill, P.H. Zaidi, R. Kumar, S.K. Nair, et al. 2021. Genome wide association mapping for heat tolerance in sub-tropical maize. BMC Genomics 22(1): 154.

López-Hernández, F., and A.J. Cortés. 2019. Last-Generation Genome–Environment Associations Reveal the Genetic Basis of Heat Tolerance in Common Bean (Phaseolus vulgaris L.). Front. Genet. 10. doi: 10.3389/fgene.2019.00954.

Lozano-Isla, F. 2021. inti: Tools and statistical procedures in plant science. R Package Version 0.1.

Ludwig, L.J., and D.T. Canvin. 1971. The Rate of Photorespiration during Photosynthesis and the Relationship of the Substrate of Light Respiration to the Products of Photosynthesis in Sunflower Leaves. Plant Physiol. 48(6): 712–719.

Lund, R.E. 1975. Tables for An Approximate Test for Outliers in Linear Models. Technometrics 17(4): 473–476.

Maalouf, F., L. Abou-Khater, Z. Babiker, A. Jighly, A.M. Alsamman, et al. 2022. Genetic Dissection of Heat Stress Tolerance in Faba Bean (Vicia faba L.) Using GWAS. Plants 11(9). doi: 10.3390/plants11091108.

Mamidi, S., S. Chikara, R.J. Goos, D.L. Hyten, D. Annam, et al. 2011. Genome-wide association analysis identifies candidate genes associated with iron deficiency chlorosis in soybean. Plant Genome 4(3): 154–164.

Maulana, F., H. Ayalew, J.D. Anderson, T.T. Kumssa, W. Huang, et al. 2018. Genome-Wide Association Mapping of Seedling Heat Tolerance in Winter Wheat. Front. Plant Sci. 9: 1272.

Melo, B.P., O.T. Fraga, J.C.F. Silva, D.O. Ferreira, O.J.B. Brustolini, et al. 2018. Revisiting the Soybean GmNAC Superfamily. Front. Plant Sci. 9: 1864.

Moellers, T.C., A. Singh, J. Zhang, J. Brungardt, M. Kabbage, et al. 2017. Main and epistatic loci studies in soybean for Sclerotinia sclerotiorum resistance reveal multiple modes of resistance in multi-environments. Sci. Rep. 7(1): 3554.

Moore, C.E., K. Meacham-Hensold, P. Lemonnier, R.A. Slattery, C. Benjamin, et al. 2021. The effect of increasing temperature on crop photosynthesis: from enzymes to ecosystems. J. Exp. Bot. 72(8): 2822–2844.

Mousavi-Derazmahalleh, M., P.E. Bayer, J.K. Hane, B. Valliyodan, H.T. Nguyen, et al. 2019. Adapting legume crops to climate change using genomic approaches. Plant Cell Environ. 42(1): 6–19.

Nagasubramanian, K., S. Jones, S. Sarkar, A.K. Singh, A. Singh, et al. 2018. Hyperspectral band selection using genetic algorithm and support vector machines for early identification of charcoal rot disease in soybean stems. Plant Methods 14: 86.

OECD, and FAO - United Nations. 2023. Oilseeds and oilseed products. OECD-FAO Agricultural Outlook. OECD

Oerke, E.-C. 2006. Crop losses to pests. J. Agric. Sci. 144(1): 31–43.

Oliveira, M.F., R.L. Nelson, I.O. Geraldi, C.D. Cruz, and J.F.F. de Toledo. 2010. Establishing a soybean germplasm core collection. Field Crops Res. 119(2): 277–289.

Paul, P.J., S. Samineni, M. Thudi, S.B. Sajja, A. Rathore, et al. 2018. Molecular Mapping of QTLs for Heat Tolerance in Chickpea. Int. J. Mol. Sci. 19(8). doi: 10.3390/ijms19082166.

Peirone, L.S., G.A. Pereyra Irujo, A. Bolton, I. Erreguerena, and L.A.N. Aguirrezábal. 2018. Assessing the Efficiency of Phenotyping Early Traits in a Greenhouse Automated Platform for Predicting Drought Tolerance of Soybean in the Field. Front. Plant Sci. 9: 587.

Poudel, S., B. Adhikari, J. Dhillon, K.R. Reddy, S.R. Stetina, et al. 2023. Quantifying the physiological, yield, and quality plasticity of Southern USA soybeans under heat stress. Plant Stress 9: 100195.

Prasad, P.V.V., R. Bheemanahalli, and S.V.K. Jagadish. 2017. Field crops and the fear of heat stress—Opportunities, challenges and future directions. Field Crops Res. 200: 114–121.

Rairdin, A., F. Fotouhi, J. Zhang, D.S. Mueller, B. Ganapathysubramanian, et al. 2022. Deep learning-based phenotyping for genome wide association studies of sudden death syndrome in soybean. Front. Plant Sci. 13: 966244.

Ruiz-Vera, U.M., M. Siebers, S.B. Gray, D.W. Drag, D.M. Rosenthal, et al. 2013. Global warming can negate the expected CO2 stimulation in photosynthesis and productivity for soybean grown in the Midwestern United States. Plant Physiol. 162(1): 410–423.

Sedivy, E.J., F. Wu, and Y. Hanzawa. 2017. Soybean domestication: the origin, genetic architecture and molecular bases. New Phytol. 214(2): 539–553.

Seetharam, K., P.H. Kuchanur, K.B. Koirala, M.P. Tripathi, A. Patil, et al. 2021. Genomic regions associated with heat stress tolerance in tropical maize (Zea mays L.). Sci. Rep. 11(1): 13730.

Sharma, V., S.S. Gangurde, S.N. Nayak, A.S. Gowda, B.S. Sukanth, et al. 2023. Genetic mapping identified three hotspot genomic regions and candidate genes controlling heat tolerance-related traits in groundnut. Front. Plant Sci. 14: 1182867.

Siebers, M.H., C.R. Yendrek, D. Drag, A.M. Locke, L. Rios Acosta, et al. 2015. Heat waves imposed during early pod development in soybean (Glycine max) cause significant yield loss despite a rapid recovery from oxidative stress. Glob. Chang. Biol. 21(8): 3114–3125.

Singh, D.P., A.K. Singh, and A. Singh. 2021. Plant Breeding and Cultivar Development. Elsevier Science.

Soltani, A., S.M. Weraduwage, T.D. Sharkey, and D.B. Lowry. 2019. Elevated temperatures cause loss of seed set in common bean (Phaseolus vulgaris L.) potentially through the disruption of source-sink relationships. BMC Genomics 20(1): 312.

Song, Q., D.L. Hyten, G. Jia, C.V. Quigley, E.W. Fickus, et al. 2015. Fingerprinting Soybean Germplasm and Its Utility in Genomic Research. G3 5(10): 1999–2006.

Song, Q., L. Yan, C. Quigley, B.D. Jordan, E. Fickus, et al. 2017a. Genetic Characterization of the Soybean Nested Association Mapping Population. Plant Genome 10(2). doi: 10.3835/plantgenome2016.10.0109.

Song, K., W.C. Yim, and B.-M. Lee. 2017b. Expression of heat shock proteins by heat stress in soybean. Plant Breed. Biotechnol. 5(4): 344–353.

Steketee, C.J., W.T. Schapaugh, T.E. Carter Jr, and Z. Li. 2020. Genome-Wide Association Analyses Reveal Genomic Regions Controlling Canopy Wilting in Soybean. G3 10(4): 1413–1425.

Su, H.-G., X.-H. Zhang, T.-T. Wang, W.-L. Wei, Y.-X. Wang, et al. 2020. Genome-Wide Identification, Evolution, and Expression of GDSL-Type Esterase/Lipase Gene Family in Soybean. Front. Plant Sci. 11: 726.

Sun, Q., C. Miao, M. Hanel, A.G.L. Borthwick, Q. Duan, et al. 2019. Global heat stress on health, wildfires, and agricultural crops under different levels of climate warming. Environ. Int. 128: 125–136.

Tacarindua, C.R.P., T. Shiraiwa, K. Homma, E. Kumagai, and R. Sameshima. 2013. The effects of increased temperature on crop growth and yield of soybean grown in a temperature gradient chamber. Field Crops Res. 154: 74–81.

Tafesse, E.G., K.K. Gali, V.B.R. Lachagari, R. Bueckert, and T.D. Warkentin. 2020. Genome-Wide Association Mapping for Heat Stress Responsive Traits in Field Pea. Int. J. Mol. Sci. 21(6). doi: 10.3390/ijms21062043.

Teixeira, E.I., G. Fischer, H. van Velthuizen, C. Walter, and F. Ewert. 2013. Global hot-spots of heat stress on agricultural crops due to climate change. Agric. For. Meteorol. 170: 206–215.

Tesfaye, K., P.H. Zaidi, S. Gbegbelegbe, C. Boeber, D.B. Rahut, et al. 2017. Climate change impacts and potential benefits of heat-tolerant maize in South Asia. Theor. Appl. Climatol. 130(3): 959–970.

Thomey, M.L., R.A. Slattery, I.H. Köhler, C.J. Bernacchi, and D.R. Ort. 2019. Yield response of field-grown soybean exposed to heat waves under current and elevated [CO2]. Glob. Chang. Biol. 25(12): 4352–4368.

Tran, L.-S.P., T.N. Quach, S.K. Guttikonda, D.L. Aldrich, R. Kumar, et al. 2009. Molecular characterization of stress-inducible GmNAC genes in soybean. Mol. Genet. Genomics 281(6): 647–664.

Ud-Din, N., B.F. Carver, and A.C. Clutter. 1992. Genetic analysis and selection for wheat yield in drought-stressed and irrigated environments. Euphytica 62(2): 89–96.

Valdés-López, O., J. Batek, N. Gomez-Hernandez, C.T. Nguyen, M.C. Isidra-Arellano, et al. 2016. Soybean Roots Grown under Heat Stress Show Global Changes in Their Transcriptional and Proteomic Profiles. Front. Plant Sci. 7: 517.

Valluru, R., M.P. Reynolds, W.J. Davies, and S. Sukumaran. 2017. Phenotypic and genome-wide association analysis of spike ethylene in diverse wheat genotypes under heat stress. New Phytol. 214(1): 271–283.

Vu, J.C.V., L.H. Allen Jr, K.J. Boote, and G. Bowes. 1997. Effects of elevated CO2 and temperature on photosynthesis and Rubisco in rice and soybean. Plant Cell Environ. 20(1): 68–76.

Vuong, T.D., H. Sonah, C.G. Meinhardt, R. Deshmukh, S. Kadam, et al. 2015. Genetic architecture of cyst nematode resistance revealed by genome-wide association study in soybean. BMC Genomics 16: 593.

Wang, Q., G. Li, K. Zheng, X. Zhu, J. Ma, et al. 2019. The Soybean Laccase Gene Family: Evolution and Possible Roles in Plant Defense and Stem Strength Selection. Genes 10(9). doi: 10.3390/genes10090701.

Wang, L., L. Liu, Y. Ma, S. Li, S. Dong, et al. 2018. Transcriptome profilling analysis characterized the gene expression patterns responded to combined drought and heat stresses in soybean. Comput. Biol. Chem. 77: 413–429.

Wang, N., H. Wang, A. Zhang, Y. Liu, D. Yu, et al. 2020. Genomic prediction across years in a maize doubled haploid breeding program to accelerate early-stage testcross testing. Theor. Appl. Genet. 133(10): 2869–2879.

Wei, Z., Q. Yuan, H. Lin, X. Li, C. Zhang, et al. 2021. Linkage analysis, GWAS, transcriptome analysis to identify candidate genes for rice seedlings in response to high temperature stress. BMC Plant Biol. 21(1): 85.

Wen, Z., R. Tan, J. Yuan, C. Bales, W. Du, et al. 2014. Genome-wide association mapping of quantitative resistance to sudden death syndrome in soybean. BMC Genomics 15(1): 809.

Wu, C., L.A. Mozzoni, D. Moseley, W. Hummer, H. Ye, et al. 2019. Genome-wide association mapping of flooding tolerance in soybean. Mol. Breed. 40(1): 4.

Xu, J., N. Driedonks, M.J.M. Rutten, W.H. Vriezen, G.-J. de Boer, et al. 2017. Mapping quantitative trait loci for heat tolerance of reproductive traits in tomato (Solanum lycopersicum). Mol. Breed. 37(5): 58.

Yang, M., and G. Wang. 2023. Heat stress to jeopardize crop production in the US Corn Belt based on downscaled CMIP5 projections. Agric. Syst. 211: 103746.

Yang, Y., C. Zhang, D. Zhu, H. He, Z. Wei, et al. 2022. Identifying candidate genes and patterns of heat-stress response in rice using a genome-wide association study and transcriptome analyses. The Crop Journal 10(6): 1633–1643.

Ye, C., M.A. Argayoso, E.D. Redoña, S.N. Sierra, M.A. Laza, et al. 2012. Mapping QTL for heat tolerance at flowering stage in rice using SNP markers. Plant Breed. 131(1): 33–41.

Young, A.J. 1991. The photoprotective role of carotenoids in higher plants. Physiol. Plant. 83(4): 702–708.

Yu, J., L. Chen, M. Xu, and B. Huang. 2012. Effects of elevated CO_2_ on physiological responses of tall fescue to elevated temperature, drought stress, and the combined stresses. Crop Sci. 52(4): 1848–1858.

Yu, X., X. Li, T. Guo, C. Zhu, Y. Wu, et al. 2016. Genomic prediction contributing to a promising global strategy to turbocharge gene banks. Nat Plants 2: 16150.

Yuan, Y., J.E. Cairns, R. Babu, M. Gowda, D. Makumbi, et al. 2019. Genome-Wide Association Mapping and Genomic Prediction Analyses Reveal the Genetic Architecture of Grain Yield and Flowering Time Under Drought and Heat Stress Conditions in Maize. Front. Plant Sci. 9. doi: 10.3389/fpls.2018.01919.

Zhang, J., H.S. Naik, T. Assefa, S. Sarkar, R.V.C. Reddy, et al. 2017a. Computer vision and machine learning for robust phenotyping in genome-wide studies. Sci. Rep. 7: 44048.

Zhang, J., A. Singh, D.S. Mueller, and A.K. Singh. 2015. Genome-wide association and epistasis studies unravel the genetic architecture of sudden death syndrome resistance in soybean. Plant J. 84(6): 1124–1136.

Zhang, Y., H. Tian, D. Chen, H. Zhang, M. Sun, et al. 2023. Cysteine-rich receptor-like protein kinases: emerging regulators of plant stress responses. Trends Plant Sci. 28(7): 776–794.

Zhang, J., X. Wang, Y. Lu, S.J. Bhusal, Q. Song, et al. 2018. Genome-wide Scan for Seed Composition Provides Insights into Soybean Quality Improvement and the Impacts of Domestication and Breeding. Mol. Plant 11(3): 460–472.

Zhang, J., Z. Wen, W. Li, Y. Zhang, L. Zhang, et al. 2017b. Genome-wide association study for soybean cyst nematode resistance in Chinese elite soybean cultivars. Mol. Breed. 37(5): 60.

Zhou, J., H. Mou, J. Zhou, M.L. Ali, H. Ye, et al. 2021. Qualification of Soybean Responses to Flooding Stress Using UAV-Based Imagery and Deep Learning. Plant Phenomics 2021: 9892570.

Zhou, X., S. Rao, E. Wrightstone, T. Sun, A.C.W. Lui, et al. 2022. Phytoene Synthase: The Key Rate-Limiting Enzyme of Carotenoid Biosynthesis in Plants. Front. Plant Sci. 13: 884720.

Zhou, R., X. Yu, C.-O. Ottosen, E. Rosenqvist, L. Zhao, et al. 2017. Drought stress had a predominant effect over heat stress on three tomato cultivars subjected to combined stress. BMC Plant Biol. 17(1): 24.

Zhou, J., J. Zhou, H. Ye, M.L. Ali, H.T. Nguyen, et al. 2020. Classification of soybean leaf wilting due to drought stress using UAV-based imagery. Comput. Electron. Agric. 175: 105576.

