## Supplementary for "Genetic Dissection of Heat Stress Tolerance in Soybean through Genome-Wide Association Studies and the Use of Genomic Prediction to Enhance Breeding Applications"

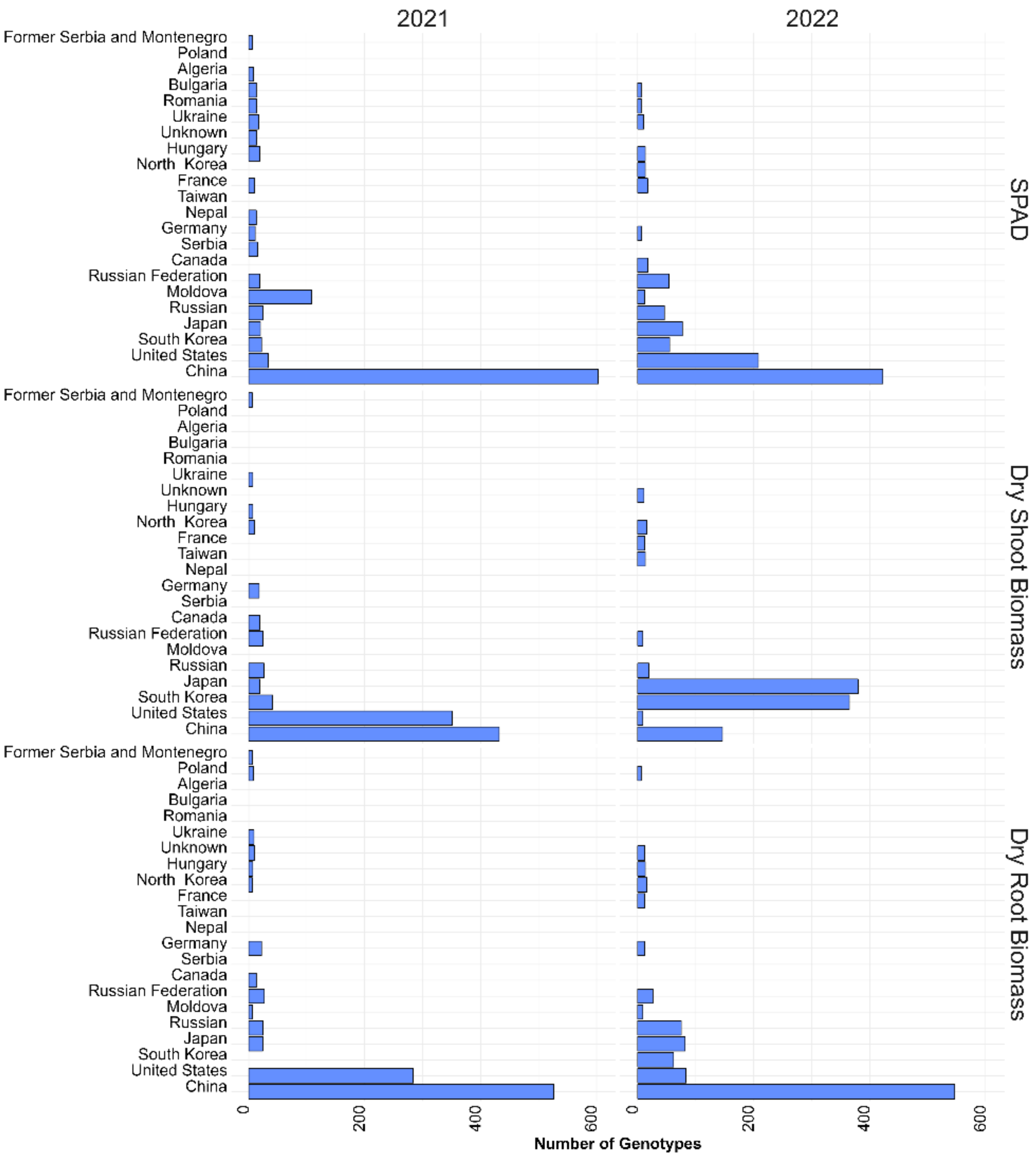


Figure S1. Country of origin of the 5% most resistant accessions for heat related traits from the scanning the USDA soybean germplasm collection with the SoySNP50K markers.

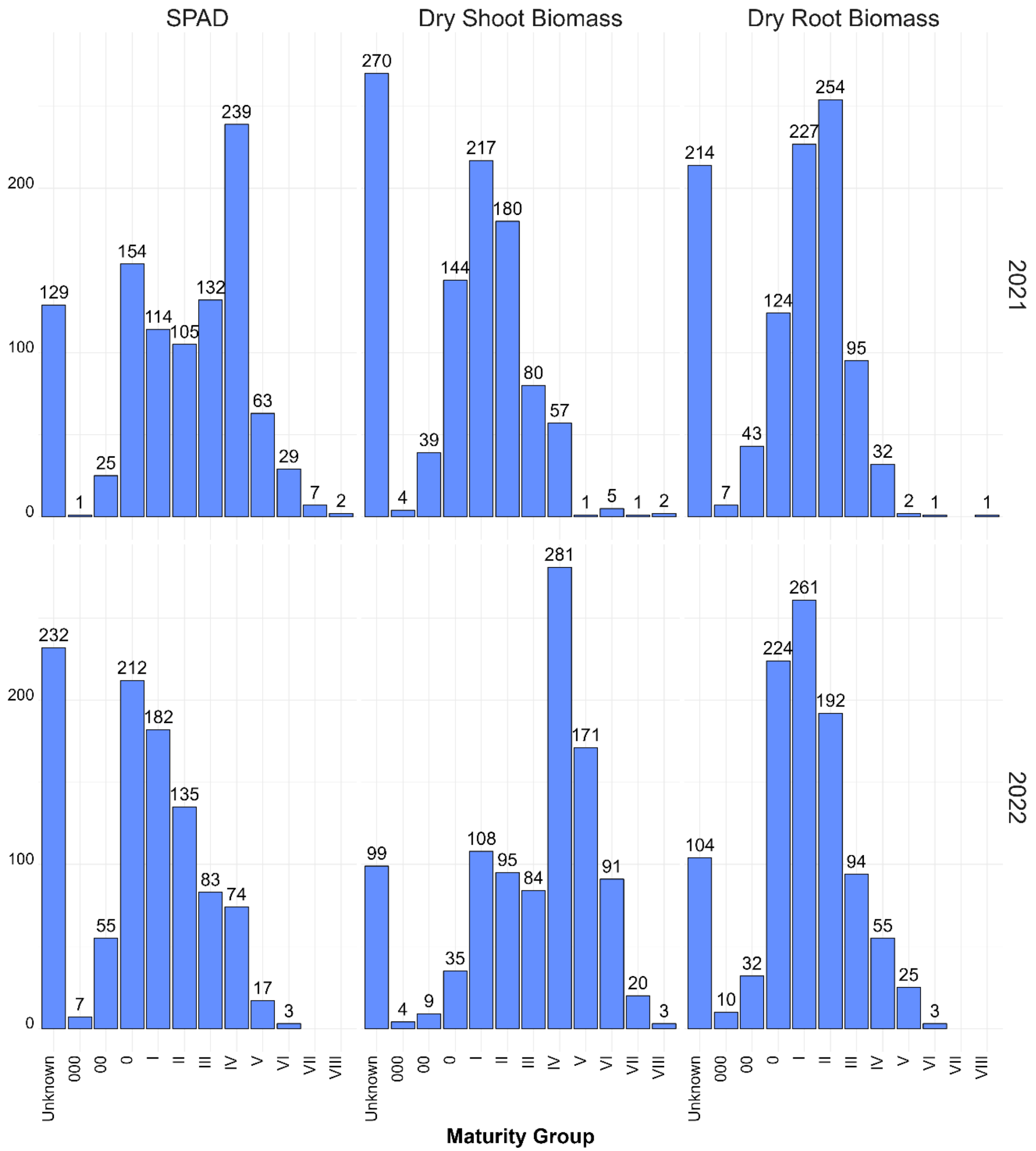


Figure S2. Maturity group distribution of the 5% most tolerant accessions for heat related traits from the entire USDA soybean germplasm collection.

**Table S1. Description of accessions, country of origin, genetic background, and maturity group used in this study.**

| Accession | Origin | Group | MG | Accession | Origin | Group | MG |
|---|---|---|---|---|---|---|---|
| 4J105-3-4 | USA | SoyNAM | III | PI257432 | Germany | PI | 0 |
| 5M20-2-5-2 | USA | SoyNAM | III | PI257436 | Germany | PI | 0 |
| CL0J095-4-6 | USA | SoyNAM | III | PI261466 | Japan | PI | III |
| CL0J173-6-8 | USA | SoyNAM | III | PI261474 | China | PI | II |
| FC30684 | China | Elite | 0 | PI266806A | China | PI | II |
| FC30692 | China | Elite | 0 | PI290116A | Hungary | PI | 0 |
| HS6-3976 | USA | SoyNAM | III | PI291319B | China | PI | 0 |
| IA1022 | USA | Elite | I | PI297513 | Russia | PI | 0 |
| IA2102 | USA | Elite | II | PI297523 | China | PI | 0 |
| IA3023 | USA | SoyNAM | III | PI297532 | China | PI | 0 |
| IA3024 | USA | Elite | III | PI323586B | Portugal | PI | II |
| IA4005 | USA | Elite | IV | PI339868E | South Korea | PI | III |
| LD01-5907 | USA | SoyNAM | III | PI347549 | Kazakhstan | PI | 0 |
| LD02-4485 | USA | SoyNAM | II | PI358316C | Japan | PI | 0 |
| LD07-3395Bf | USA | Elite | IV | PI358323 | China | PI | 0 |
| LG03-2979 | USA | SoyNAM | III | PI361058 | Germany | PI | 0 |
| LG04-4717 | USA | SoyNAM | III | PI361101 | South Korea | PI | III |
| LG05-4464 | USA | SoyNAM | III | PI378658 | Ukraine | PI | 0 |
| LG05-4832 | USA | SoyNAM | III | PI378674A | Bulgaria | PI | 0 |
| LG90-2550 | USA | SoyNAM | III | PI379559D | Japan | PI | III |
| MN1410 | USA | Elite | I | PI379561 | Japan | PI | III |
| NE3001 | USA | SoyNAM | III | PI391577 | China | PI | II |
| PI153251 | Unknown | PI | 0 | PI391586 | China | PI | III |
| PI153280 | France | PI | II | PI398813 | South Korea | PI | III |
| PI154196 | Netherlands | PI | 0 | PI398881 | South Korea | SoyNAM | III |
| PI157421 | South Korea | PI | III | PI404160B | Georgia | PI | III |
| PI167240 | Turkey | PI | III | PI404166 | China | PI | III |
| PI171450 | Japan | PI | III | PI404169B | China | PI | III |
| PI173994 | South Korea | PI | III | PI404188A | China | SoyNAM | II |
| PI189903 | France | PI | 0 | PI407653 | China | PI | III |
| PI189916 | China | PI | I | PI407656 | China | PI | II |
| PI189930 | France | PI | II | PI407659A | China | PI | II |
| PI189961 | France | PI | 0 | PI407746 | China | PI | III |
| PI189969 | France | PI | III | PI407810 | South Korea | PI | III |
| PI196149 | Japan | PI | III | PI416773 | Japan | PI | II |
| PI200548 | Japan | PI | III | PI416868A | Japan | PI | III |
| PI227325 | Japan | PI | I | PI417091 | Japan | PI | II |
| PI232996 | Germany | PI | 0 | PI417198 | Japan | PI | III |
| PI238921 | Germany | PI | 0 | PI417458 | Japan | PI | 0 |
| PI248403 | Serbia | PI | 0 | PI417513B | Eastern Europe | PI | I |
| PI253650A | China | PI | II | PI417522 | Croatia | PI | 0 |
| PI253651C | China | PI | III | PI417552 | Poland | PI | 0 |
| PI253653D | China | PI | I | PI417559 | Poland | PI | III |
| PI253658A | China | PI | I | PI424148 | South Korea | PI | 0 |
| PI253660B | China | PI | III | PI427136 | South Korea | SoyNAM | III |

| Accession | Origin | Group | MG | Accession | Origin | Group | MG |
|---|---|---|---|---|---|---|---|
| PI430597 | China | PI | II | PI437757 | China | PI | I |
| PI430619 | China | PI | III | PI437786 | China | PI | I |
| PI437090 | Russia | PI | 0 | PI437803 | China | PI | II |
| PI437091 | Russia | PI | I | PI437812 | China | PI | 0 |
| PI437098 | Russia | PI | I | PI437840A | China | PI | II |
| PI437100 | Russia | PI | 0 | PI437846 | China | PI | I |
| PI437121B | Russia | PI | II | PI437923 | China | PI | 0 |
| PI437122 | Russia | PI | II | PI437949 | China | PI | I |
| PI437124 | Georgia | PI | III | PI437950 | China | PI | II |
| PI437145B | Russia | PI | II | PI437973 | China | PI | II |
| PI437156C | Russia | PI | I | PI437982 | China | PI | 0 |
| PI437169B | Russia | SoyNAM | II | PI437995A | China | PI | 0 |
| PI437174B | Russia | PI | I | PI438031 | China | PI | I |
| PI437202 | Moldova | PI | 0 | PI438094B | China | PI | I |
| PI437207 | Moldova | PI | 0 | PI438103 | China | PI | II |
| PI437230 | Moldova | PI | 0 | PI438133B | China | PI | II |
| PI437238 | Moldova | PI | 0 | PI438139 | China | PI | II |
| PI437255 | Moldova | PI | 0 | PI438148 | China | PI | 0 |
| PI437263 | Moldova | PI | 0 | PI438173 | China | PI | II |
| PI437306A | Russia | PI | 0 | PI438194 | China | PI | II |
| PI437340B | Russia | PI | II | PI438218 | China | PI | I |
| PI437343 | Russia | PI | I | PI438239A | China | PI | 0 |
| PI437356 | Russia | PI | II | PI438259B | China | PI | III |
| PI437377 | Russia | PI | III | PI438292 | Japan | PI | I |
| PI437399 | Russia | PI | II | PI438376 | France | PI | I |
| PI437425 | Russia | PI | I | PI438434 | Morocco | PI | II |
| PI437427B | Russia | PI | II | PI438469 | Romania | PI | 0 |
| PI437462A | Russia | PI | II | PI438503A | USA | PI | II |
| PI437477A | Russia | PI | I | PI445819 | Germany | PI | I |
| PI437509 | Russia | PI | I | PI445827B | Romania | PI | 0 |
| PI437519 | Russia | PI | I | PI445833 | Romania | PI | 0 |
| PI437553 | China | PI | 0 | PI445845 | China | PI | III |
| PI437558 | China | PI | I | PI458052 | South Korea | PI | III |
| PI437561 | China | PI | 0 | PI458061A | South Korea | PI | III |
| PI437581 | China | PI | II | PI458110 | South Korea | PI | III |
| PI437585 | China | PI | II | PI458307A | South Korea | PI | III |
| PI437592 | China | PI | II | PI458506 | China | PI | II |
| PI437594A | China | PI | I | PI458507 | China | PI | III |
| PI437651B | China | PI | II | PI458517 | China | PI | III |
| PI437674 | China | PI | III | PI458519A | China | PI | II |
| PI437690 | China | PI | III | PI458520 | China | PI | II |
| PI437712 | China | PI | 0 | PI458521 | China | PI | III |
| PI437715 | China | PI | II | PI458522 | China | PI | II |
| PI437716A | China | PI | I | PI458825B | China | PI | I |
| PI437753A | China | PI | 0 | PI461509 | China | PI | I |

| Accession | Origin | Group | MG | Accession | Origin | Group | MG |
|---|---|---|---|---|---|---|---|
| PI464877 | China | PI | III | PI506887 | Japan | PI | III |
| PI464878 | China | PI | II | PI507171 | Japan | PI | III |
| PI464880 | China | PI | II | PI507195 | Japan | PI | II |
| PI464884 | China | PI | II | PI507201 | Japan | PI | 0 |
| PI464914B | China | PI | III | PI507487 | Japan | PI | III |
| PI464915A | China | PI | II | PI507491 | Japan | PI | III |
| PI467307 | China | PI | I | PI507685B | Ukraine | PI | 0 |
| PI467310 | China | PI | II | PI507688 | Moldova | PI | 0 |
| PI467311A | China | PI | I | PI507717 | North Korea | PI | I |
| PI467312 | China | PI | II | PI512322C | Georgia | PI | I |
| PI467313 | China | PI | 0 | PI518706A | China | PI | I |
| PI467324 | China | PI | I | PI518751 | Former Serbia and Montenegro | SoyNAM | II |
| PI467327 | China | PI | II | PI518757 | Taiwan | PI | III |
| PI467328 | China | PI | I | PI522188A | Russia | PI | I |
| PI467332 | China | PI | II | PI524994 | Russia | PI | I |
| PI468381 | Japan | PI | II | PI532462A | China | PI | III |
| PI468384 | China | PI | III | PI538377 | China | PI | III |
| PI468385 | China | PI | III | PI538393 | China | PI | I |
| PI468914 | China | PI | III | PI538400 | China | PI | II |
| PI470223 | China | PI | II | PI538403 | Japan | PI | I |
| PI470227B | China | PI | III | PI538410B | Japan | PI | I |
| PI471899 | Indonesia | PI | III | PI540739 | China | PI | I |
| PI475810 | China | PI | II | PI54591 | China | PI | III |
| PI475811B | China | PI | II | PI54608-1 | China | PI | II |
| PI475818 | China | PI | III | PI548316 | China | PI | III |
| PI475820 | China | PI | II | PI548329 | Japan | PI | I |
| PI475822B | China | PI | III | PI548336 | Russia | PI | I |
| PI476344 | Uzbekistan | PI | II | PI548349 | North Korea | PI | III |
| PI476345 | Moldova | PI | I | PI548373 | China | PI | III |
| PI476348 | Ukraine | PI | I | PI54854 | China | PI | I |
| PI476911 | Vietnam | PI | II | PI549021A | China | PI | III |
| PI479711 | China | PI | II | PI549031 | China | PI | III |
| PI479713 | China | PI | II | PI561227 | China | PI | II |
| PI479718B | China | PI | II | PI561230 | China | PI | II |
| PI479719 | China | PI | I | PI561232 | China | PI | I |
| PI479724A | China | PI | II | PI561242 | China | PI | 0 |
| PI479729 | China | PI | III | PI561315 | China | PI | I |
| PI479738 | China | PI | II | PI561333 | China | PI | I |
| PI479740 | China | PI | III | PI561346 | China | PI | I |
| PI504485 | Japan | PI | I | PI561349 | China | PI | II |
| PI504490 | Taiwan | PI | II | PI561370 | China | SoyNAM | III |
| PI504497 | Taiwan | PI | II | PI561377 | Japan | PI | II |
| PI506527 | Japan | PI | III | PI567154 | Japan | PI | II |
| PI506528 | Japan | PI | III | PI567159A | China | PI | I |
| PI506529 | Japan | PI | III | PI567161 | China | PI | II |

| Accession | Origin | Group | MG | Accession | Origin | Group | MG |
|---|---|---|---|---|---|---|---|
| PI567170A | China | PI | II | PI578439 | Vietnam | PI | III |
| PI567170B | China | PI | II | PI578473A | China | PI | III |
| PI567212B | Russia | PI | 0 | PI578474 | China | PI | I |
| PI567214B | Russia | PI | I | PI578485A | China | PI | 0 |
| PI567217A | Russia | PI | 0 | PI578499A | China | PI | II |
| PI567225 | Moldova | PI | 0 | PI578499B | China | PI | II |
| PI567229A | Russia | PI | I | PI588008A | China | PI | III |
| PI567241 | China | PI | II | PI592907C | Russia | PI | I |
| PI567250B | China | PI | I | PI592908 | Russia | PI | II |
| PI567255A | China | PI | I | PI592910 | Russia | PI | II |
| PI567261B | China | PI | II | PI592911B | Russia | PI | I |
| PI567262D | China | PI | II | PI592912A | Russia | PI | I |
| PI567264A | China | PI | II | PI592913 | Russia | PI | II |
| PI567266A | China | PI | II | PI594394 | China | PI | III |
| PI567267A | China | PI | II | PI594898 | China | PI | I |
| PI567275 | Japan | PI | II | PI594902 | China | PI | I |
| PI567277 | Japan | PI | II | PI597391C | Ukraine | PI | 0 |
| PI567278 | Japan | PI | II | PI597397A | Russia | PI | I |
| PI567351A | China | PI | II | PI597400 | Russia | PI | 0 |
| PI567365 | China | PI | III | PI597405B | Ukraine | PI | I |
| PI567417B | China | PI | I | PI597467 | China | PI | 0 |
| PI567538B | China | PI | II | PI597482 | South Korea | PI | III |
| PI567583A | China | PI | III | PI597651 | China | PI | 0 |
| PI567595A | China | PI | III | PI598124 | USA | PI | III |
| PI567619 | China | PI | III | PI602497A | China | PI | I |
| PI567644 | China | PI | III | PI603151A | North Korea | PI | I |
| PI567729 | China | PI | III | PI603297 | China | PI | 0 |
| PI567774B | China | PI | III | PI603306 | China | PI | 0 |
| PI574480B | China | PI | III | PI603334 | China | PI | I |
| PI574486 | China | SoyNAM | III | PI603335B | China | PI | II |
| PI578360 | China | PI | II | PI603337A | China | PI | I |
| PI578362 | China | PI | I | PI603339A | China | PI | I |
| PI578363 | China | PI | II | PI603367 | China | PI | I |
| PI578364 | China | PI | II | PI603371 | China | PI | I |
| PI578366 | China | PI | III | PI603412B | China | PI | II |
| PI578367 | China | PI | III | PI603422B | China | PI | II |
| PI578374 | China | PI | I | PI603424C | China | PI | I |
| PI578375B | China | PI | I | PI603426F | China | PI | I |
| PI578376 | China | PI | II | PI603428D | China | PI | III |
| PI578380A | China | PI | I | PI603429A | China | PI | 0 |
| PI578382 | China | PI | I | PI603438E | China | PI | III |
| PI578384 | China | PI | I | PI603442 | China | PI | III |
| PI578385 | China | PI | I | PI603444A | China | PI | II |
| PI578386 | China | PI | I | PI603452 | China | PI | III |
| PI578416 | China | PI | II | PI603470 | China | PI | II |

| Accession | Origin | Group | MG | Accession | Origin | Group | MG |
|---|---|---|---|---|---|---|---|
| PI603546A | China | PI | I | PI84921 | North Korea | PI | II |
| PI603560 | China | PI | III | PI85009-1 | Japan | PI | III |
| PI603594 | China | PI | II | PI85356 | South Korea | PI | III |
| PI603596 | China | PI | III | PI86006 | Japan | PI | III |
| PI603655 | China | PI | III | PI86145 | Japan | PI | III |
| PI603660 | China | PI | II | PI86449 | Japan | PI | III |
| PI603662B | China | PI | II | PI87600-1 | North Korea | PI | III |
| PI603674 | China | PI | III | PI87618 | North Korea | PI | III |
| PI603704A | China | PI | I | PI87631-1 | Japan | PI | III |
| PI603712 | China | PI | 0 | PI87634 | Japan | PI | III |
| PI603747 | China | PI | II | PI88289 | China | PI | III |
| PI603749 | China | PI | II | PI88292 | China | PI | III |
| PI603912 | North Korea | PI | III | PI88294-1 | China | PI | II |
| PI603915C | North Korea | PI | III | PI88295 | China | PI | I |
| PI612611 | North Korea | PI | III | PI88305 | China | PI | III |
| PI612711B | China | PI | I | PI88306 | China | PI | III |
| PI62202 | China | PI | III | PI88788 | China | PI | III |
| PI631398 | USA | PI | III | PI89003-1 | China | PI | II |
| PI633730 | USA | PI | II | PI89008 | China | PI | II |
| PI633731 | USA | PI | II | PI89060 | China | PI | I |
| PI639283 | USA | PI | II | PI89134 | North Korea | PI | III |
| PI639285 | USA | PI | III | PI89152 | North Korea | PI | III |
| PI639693 | USA | PI | II | PI89153 | North Korea | PI | II |
| PI643146 | USA | PI | IV | PI89154 | North Korea | PI | II |
| PI660989 | USA | PI | III | PI89156 | North Korea | PI | II |
| PI68685 | China | PI | II | PI89773 | China | PI | III |
| PI68722 | China | PI | 0 | PI90392 | China | PI | III |
| PI68788 | China | PI | II | PI91091 | China | PI | II |
| PI68815 | China | PI | II | PI91102 | China | PI | II |
| PI70241 | China | PI | I | PI91120-3 | China | PI | III |
| PI70463 | China | PI | II | PI91162 | China | PI | III |
| PI71161 | China | PI | I | PI91341 | China | PI | III |
| PI72232 | China | PI | III | PI91349 | China | PI | III |
| PI79586 | China | PI | II | PI91559 | China | PI | II |
| PI79691-4 | China | PI | III | PI91725-3 | North Korea | PI | II |
| PI79756 | China | PI | II | PI92603 | China | PI | II |
| PI79870-1 | China | PI | I | PI92611 | China | PI | II |
| PI80459 | Japan | PI | III | PI92683 | China | PI | II |
| PI80461 | Japan | PI | III | PI96162 | North Korea | PI | III |
| PI80469 | Japan | PI | II | PI96199 | China | PI | III |
| PI80831 | China | PI | III | PI96322 | North Korea | PI | III |
| PI81044-2 | Japan | PI | III | PI96786-1 | North Korea | PI | III |
| PI81667 | China | PI | III | U03-100612 | USA | SoyNAM | I |
| PI82278 | South Korea | PI | III | U11-917032 | USA | Elite | I |
| PI84611 | South Korea | PI | III | U11-920017 | USA | Elite | II |

**Table S2. Dates for planting and phenotyping for each treatment and replication in 2021 and 2022. HT is the heat treatment and OT is the optimal treatment (control). Replications were staggered to allow for complete phenotyping of a replication in each day.**

| Planting Dates | | | |
|---|---|---|---|
| 2021 | | 2022 | |
| July 29 | HT Replication 1 | July 25 | HT Replication 1 |
| July 30 | OT Replication 1 | July 26 | OT Replication 1 |
| August 2 | HT Replication 2 | July 28 | HT Replication 2 |
| August3 | OT Replication 2 | July 29 | OT Replication 2 |
| August 5 | HT Replication 3 | August 1 | HT Replication 3 |
| August 6 | OT Replication 3 | August 2 | OT Replication 3 |
| August 9 | HT Replication 4 | August 4 | HT Replication 4 |
| August 10 | OT Replication 4 | August 5 | OT Replication 4 |
| **Phenotyping Dates** | | | |
| **2021** | | **2022** | |
| August 26 | HT Replication 1 | August 22 | HT Replication 1 |
| August 27 | OT Replication 1 | August 23 | OT Replication 1 |
| August 30 | HT Replication 2 | August 25 | HT Replication 2 |
| August 31 | OT Replication 2 | August 26 | OT Replication 2 |
| September 2 | HT Replication 3 | August 29 | HT Replication 3 |
| September 3 | OT Replication 3 | August 30 | OT Replication 3 |
| September 6 | HT Replication 4 | September 1 | HT Replication 4 |
| September 7 | OT Replication 4 | September 2 | OT Replication 4 |

**Table S3. ANOVA results of fixed effects for mixed linear models with the phenotypic traits as the response variables.**

| Trait | Year | Source of Variation | F-Value | df |
|---|---|---|---|---|
| SPAD | 2021 | Genotype (G) | 1.69*** | 411 |
| | | Treatment (T) | 0.45 | 1 |
| | | Replication (R) | 0.03 | 1 |
| | | Genotype x Treatment (GT) | 0.91 | 411 |
| | 2022 | Genotype (G) | 2.66*** | 446 |
| | | Treatment (T) | 7.27** | 1 |
| | | Replication (R) | 0.05 | 1 |
| | | Genotype x Treatment (GT) | 1.41*** | 446 |
| Canopy Temperature | 2021 | Genotype (G) | 1.07 | 411 |
| | | Treatment (T) | 0.78 | 1 |
| | | Replication (R) | 6.04* | 1 |
| | | Genotype x Treatment (GT) | 1.19** | 398 |
| | 2022 | Genotype (G) | 1.09 | 446 |
| | | Treatment (T) | 62.59*** | 1 |
| | | Replication (R) | 0.37 | 1 |
| | | Genotype x Treatment (GT) | 1.13* | 443 |
| Fresh Shoot Biomass | 2022 | Genotype (G) | 3.87*** | 446 |
| | | Treatment (T) | 12.80** | 1 |
| | | Replication (R) | 2.52 | 1 |
| | | Genotype x Treatment (GT) | 1.44*** | 446 |
| Dry Shoot Biomass | 2021 | Genotype (G) | 2.46*** | 411 |
| | | Treatment (T) | 0.29 | 1 |
| | | Replication (R) | 9.21** | 1 |
| | | Genotype x Treatment (GT) | 1.15** | 383 |
| | 2022 | Genotype (G) | 3.68*** | 446 |
| | | Treatment (T) | 41.30*** | 1 |
| | | Replication (R) | 0.02 | 1 |
| | | Genotype x Treatment (GT) | 1.46*** | 446 |
| Fresh Root Biomass | 2022 | Genotype (G) | 3.28*** | 446 |
| | | Treatment (T) | 0.01 | 1 |
| | | Replication (R) | 1.34 | 1 |
| | | Genotype x Treatment (GT) | 1.31*** | 446 |
| Dry Root Biomass | 2021 | Genotype (G) | 2.10*** | 411 |
| | | Treatment (T) | 0.29 | 1 |
| | | Replication (R) | 3.83 | 1 |
| | | Genotype x Treatment (GT) | 1.21*** | 379 |
| | 2022 | Genotype (G) | 2.98*** | 446 |
| | | Treatment (T) | 0.39 | 1 |
| | | Replication (R) | 2.98 | 1 |
| | | Genotype x Treatment (GT) | 1.27*** | 446 |
| Stomatal Conductance | 2022 | Genotype (G) | 1.53*** | 446 |
| | | Treatment (T) | 7.99** | 1 |
| | | Replication (R) | 3.31 | 1 |
| | | Genotype x Treatment (GT) | 1.18*** | 446 |
| Quantum Efficiency of PSII | 2022 | Genotype (G) | 1.31*** | 446 |
| | | Treatment (T) | 3.02 | 1 |
| | | Replication (R) | 5.91* | 1 |
| | | Genotype x Treatment (GT) | 0.99 | 446 |

***, **, * Significance at 0.01 level, 0.05 level, and 0.1 level, respectively

**Table S4. Complete listing of significant SNPs detected by the SVEN methodology in heat and optimal temperature treatments.**

| SNP | Chr | Pos | Trait | Environment | MIP |
|---|---|---|---|---|---|
| *ss715580772* | 1 | 6708220 | Dry Root Biomass | Heat 2022 | 0.6961 |
| *ss715580790* | 1 | 7320074 | Fresh Shoot Biomass | Heat 2022 | 0.6037 |
| *ss715579731* | 1 | 48593077 | Fresh Shoot Biomass | Heat 2022 | 0.9951 |
| *ss715582342* | 2 | 38935468 | Quantum Efficiency | Optimal 2022 | 0.5057 |
| *ss715583256* | 2 | 45904811 | SPAD | Heat 2022 | 1.0000 |
| *ss715583466* | 2 | 47857148 | Stomatal Conductance | Optimal 2022 | 0.6309 |
| *ss715585824* | 3 | 37499390 | Canopy Temperature | Optimal 2022 | 0.7652 |
| *ss715589293* | 4 | 7667305 | Fresh Root Biomass | Optimal 2022 | 0.6358 |
| *ss715588916* | 4 | 51099840 | Fresh Shoot Biomass | Heat 2022 | 1.0000 |
| | | | Dry Root Biomass | Heat 2022 | 0.8893 |
| *ss715590067* | 5 | 3508940 | Dry Shoot Biomass | Optimal 2022 | 0.8435 |
| *ss715591790* | 5 | 40974254 | Dry Shoot Biomass | Heat 2022 | 0.7328 |
| *ss715593740* | 6 | 17401316 | Stomatal Conductance | Optimal 2022 | 0.9568 |
| *ss715602620* | 8 | 5968621 | SPAD | Optimal 2022 | 0.6815 |
| *ss715602692* | 8 | 7577565 | SPAD | Optimal 2022 | 0.6224 |
| *ss715601977* | 8 | 42381368 | Quantum Efficiency | Heat 2022 | 0.5006 |
| *ss715603810* | 9 | 39306136 | Fresh Shoot Biomass | Optimal 2022 | 0.6013 |
| *ss715603883* | 9 | 39879569 | Fresh Shoot Biomass | Optimal 2022 | 0.5546 |
| *ss715604755* | 9 | 47022602 | Fresh Shoot Biomass | Optimal 2022 | 0.7420 |
| | | | Fresh Root Biomass | Optimal 2022 | 0.7420 |
| *ss715607454* | 10 | 45093186 | Dry Shoot Biomass | $H_{tl}$ | 0.8829 |
| *ss715607988* | 10 | 49420201 | Fresh Shoot Biomass | Heat 2022 | 0.9914 |
| *ss715609383* | 11 | 26799490 | Dry Shoot Biomass | Optimal 2022 | 0.6481 |
| *ss715610072* | 11 | 29249534 | Dry Shoot Biomass | Heat 2022 | 0.9799 |
| *ss715610434* | 11 | 33180864 | Fresh Root Biomass | Optimal 2022 | 0.9635 |
| | | | Fresh Root Biomass | Optimal 2022 | 0.9877 |
| *ss715613671* | 12 | 9209526 | Dry Root Biomass | Optimal 2022 | 0.9881 |
| *ss715611949* | 12 | 23457337 | SPAD | Optimal 2022 | 0.7069 |
| *ss715612319* | 12 | 33616569 | Dry Shoot Biomass | Heat 2022 | 0.5051 |
| *ss715612326* | 12 | 33745390 | Fresh Shoot Biomass | Heat 2022 | 0.5974 |
| *ss715612610* | 12 | 35873634 | Fresh Root Biomass | Optimal 2022 | 0.9423 |
| | | | Dry Root Biomass | Optimal 2022 | 0.6369 |
| *ss715613794* | 13 | 20597010 | Quantum Efficiency | Heat 2022 | 0.6994 |
| *ss715615430* | 13 | 33543422 | Dry Root Biomass | Heat 2022 | 0.5499 |
| *ss715616040* | 13 | 38501835 | Stomatal Conductance | Optimal 2022 | 0.9970 |
| *ss715625497* | 16 | 756848 | Canopy Temperature | Heat 2022 | 0.5136 |
| *ss715628154* | 17 | 7208187 | Dry Shoot Biomass | Optimal 2022 | 0.5113 |
| *ss715627447* | 17 | 38333431 | Fresh Shoot Biomass | Heat 2022 | 0.7855 |
| | | | Dry Root Biomass | Optimal 2022 | 0.7876 |
| *ss715632400* | 18 | 56876857 | Dry Shoot Biomass | Optimal 2022 | 0.5873 |
| *ss715635407* | 19 | 45015436 | Stomatal Conductance | Heat 2022 | 0.8025 |
| *ss715637111* | 20 | 27438278 | Dry Root Biomass | Optimal 2022 | 0.5797 |

The chromosome and position are based on the Wm82.a2 reference genome construction.
MIP, marginal inclusion probability. MIP ≥ 0.05 were declared as significant SNPs.

**Table S5. Comparison of significant SNPs detected via the SVEN methodology compared with the results from the TASSEL and GAPIT:FarmCPU methodology. A subset of SNPs from SVEN that were near significant SNPs (p ≤ 0.05) detected in the other methodologies.**

| | | | TASSEL | | | | FarmCPU | | | |
|---|---|---|---|---|---|---|---|---|---|---|
| SVEN SNP | Trait | Environment | SNP | Trait | P-value | Distance | SNP | Trait | P-value | Distance |
| *ss715583256* | SPAD | Heat | *ss715583258* | SPAD | 0.049 | 6 | *ss715583256* | SPAD | 0.022 | 0 |
| *ss715625497* | Canopy Temperature | Heat | *ss715625515* | Canopy Temperature | 0.029 | 5.85 | - | - | - | - |
| *ss715585824* | Canopy Temperature | Optimal | - | - | - | - | *ss715585822* | Canopy Temperature | 0.023 | 7.28 |
| *ss715604755* | FSB<br>FRB | Optimal | *ss715604754* | FSB | 0.012 | 1.89 | - | - | - | - |
| *ss715627447* | FSB<br>DRB | Heat<br>Optimal | *ss715627444* | DRB | 0.019 | 31.99 | *ss715627439* | DRB | 0.048 | 62.18 |
| *ss715628154* | DSB | Optimal | *ss715628151* | DSB | 0.036 | 15.03 | *ss715628154* | DRB | 0.021 | 0 |
| *ss715607454* | DSB | $H_{tl}$ | *ss715607454* | DSB | <0.001 | 0 | *ss715607454* | DSB | <0.001 | 0 |
| *ss715635407* | Stomatal Conductance | Heat | *ss715635401* | Conductance | 0.041 | 74.03 | *ss715635401*<br>*ss715635408* | Conductance | 0.037<br>0.05 | 74.03<br>26.92 |
| *ss715601977* | Quantum Efficiency | Heat | - | - | - | - | *ss715601977* | Quantum | 0.001 | 0 |

The p values reported here are the values reported for the selected SNPs from TASSEL or Farm CPU. The distance is in KBP.
